# Molecular Simulation of Nonfacilitated Membrane Permeation

**DOI:** 10.1101/029140

**Authors:** Ernest Awoonor-Williams, Christopher N. Rowley

**Affiliations:** Department of Chemistry, Memorial University of Newfoundland, St. John’s, NL A1B 3X7 Canada

**Keywords:** Review, Membrane, Lipid Bilayer, Permeation, Non-facilitated, Solubility-diffusion Model, Molecular Dynamics, Diffusion, PMF, Potential of Mean Force, Polarizable, Coarse Grain

## Abstract

This is a review. Non-electrolytic compounds typically cross cell membranes by passive diffusion. The rate of permeation is dependent on the chemical properties of the solute and the composition of the lipid bilayer membrane. Predicting the permeability coefficient of a solute is important in pharmaceutical chemistry and toxicology. Molecular simulation has proven to be a valuable tool for modeling permeation of solutes through a lipid bilayer. In particular, the solubility-diffusion model has allowed for the quantitative calculation of permeability coefficients. The underlying theory and computational methods used to calculate membrane permeability are reviewed. We also discuss applications of these methods to examine the permeability of solutes and the effect of membrane composition on permeability. The application of coarse grain and polarizable models is discussed.

## 1. Introduction

One function of a biological membrane is to serve as a barrier between the cytosol and extracellular environment (1; 2; 3). Intracellular compartments like mitochondria, the nucleus, etc, are also enclosed by membranes. The primary component of these membranes is amphiphilic lipids. These lipids consist of a polar or ionic headgroup and a tail that is comprised of one or more hydrocarbon chains. The hydrophilicity of the head groups and the hydrophobicity of the tails cause the lipids to spontaneously self-assemble into planar bilayers where the headgroups face the solution and the tails form a hydrophobic interior layer.

Solutes crossing the membrane must pass through the nonpolar lipid tail region of the membrane. Compounds that are more soluble in bulk water than they are in the nonpolar membrane interior will tend not to partition into the membrane. This simple mechanism allows these thin bilayers (~ 40 – 50 Å thick) to provide an effective barrier for highly water-soluble compounds like ions and sugars. In pure lipid bilayers, the rates of permeation of these species are very low. In biological membranes, rapid permeation of these compounds can be facilitated by membrane proteins like channels or transporters (4; 5; 6).

Two distinct mechanisms have been proposed for permeation in the absence of a protein facilitator: passage through a transient water pore and direct permeation through the membrane (7). Rare fluctuations in the bilayer structure can form transient water pores that allow ionic compounds to cross the bilayer while still solvated by water. This avoids the high thermodynamic penalty for dehydrating the ion. The second mechanism applies to non-electrolytic solutes, which can permeate directly through the membrane. This review will focus on molecular simulations of this second mechanism.

Many non-electrolytic compounds can permeate directly through the membrane because there is a significant probability for them to partition into the interior of the membrane. These compounds are generally only moderately soluble in aqueous solutions due to the lack of strong electrostatic interactions with water molecules. The London dispersion attractions between these solutes and the lipid tails can be competitive to their interactions with water, so the thermodynamic penalty for these compounds to enter the interior of the membrane is attenuated or even eliminated. To varying degrees, these compounds can undergo direct permeation without facilitation by a transmembrane protein.

The rate of permeation of a solute across a membrane is defined by its flux (*J*), which gives the number of molecules that cross a unit area of the membrane per unit time (e.g., *μ*mol/s/cm^2^). The flux is the product of the concentration gradient of the solute across the membrane (Δ*C*) and the permeability coefficient (*P_m_*),

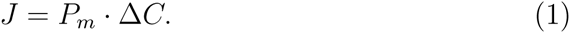

*P_m_* depends on the type of permeating solute, the membrane composition, and the conditions that permeation occurs under (e.g., temperature). It has units of distance per unit time and is commonly reported in cm/s. For a constant set of conditions and membrane composition, the permeability coefficient provides a measure of the intrinsic membrane permeability of a solute. *P_m_* is therefore the standard experimental and computational measure of the permeability of a solute.

### 1.1. Experimental determination of permeability

A variety of experimental techniques are used to determine permeability coefficients. One of the most general techniques uses a planar bilayer separating two cells. A concentration gradient is created between the two cells and the change in concentration is measured after a time interval. Several techniques have been used to measure the solution concentrations, including radiolabeling (8), UV-Vis spectroscopy (9), and LC-MS (10).

The concentration of electrochemically-active species can be measured by selective microelectrodes near the surface of the membrane. In some cases, the electrode measures the permeating species directly (e.g., the permeation of K^+^ (11)). Weak acids and bases typically permeate in their neutral, conjugate form, but their permeation can be measured by microelectrodes that detect the ionic forms that are formed in the solution at the surface of the membrane. For example, the permeability of ammonia has been measured by ammonium -selective electrodes (12). Similarly, the permeation of many weak acids has been measured by pH-sensitive microelectrodes that detect the small changes in pH at the surface of the membrane (13; 14). H_2_O_2_ permeability has been determined by an O_2_-selective electrode by measuring concentrations of O_2_ formed by the reduction of H_2_O_2_ inside the cell (15; 16).

Magnetic resonance techniques can be used to study the permeability of certain solutes. Electron paramagnetic resonance (EPR) can be used to measure the permeability of solutes with unpaired electrons, such as NO (17; 18; 19) and O_2_ (20; 21). This technique is particularly powerful because the rate of diffusion at different depths within the membrane can be measured. ^1^H and ^17^O Nuclear Magnetic Resonance (NMR) has also been used to measure the rate of water permeation. Water molecules that permeate into a liposome containing paramagnetic metal complexes (e.g., Mn(II), Ln(III), or Gd(III)) will undergo a rapid spin relaxation, allowing the rate at which water molecules cross the membrane to be determined (22).

Fluorescence spectroscopy can be used to measure the permeability of solutes that affect fluorophores loaded inside liposomes. The transmembrane permeation of protons was measured by the change in fluorescence of pyranine, a pH-sensitive dye (11). Water permeability has been calculated from the increase in self-quenching of carboxyfluorescein as the volume of the liposome decreases in response to osmotic pressure (23). The permeation of the fluorophore itself can also be measured if the fluorescence is affected by the contents of the liposome. For example, the permeation of tetracyclines was measured by the observation of the enhanced fluorescence of a Tet-repressor-bound tetracycline inside a liposome (24). The rate of permeation of aromatic-containing acids, like salicylic acid, into liposomes has been measured by monitoring the fluorescence of Tb^3+^-carboxylate complexes (25).

One limitation to permeability measurements using liposomes is that stopped-flow devices are often used to create the concentration gradient between the solution and the interior of the liposome. Pohl and coworkers noted that the mixing time associated with this technique means that they cannot be used to study very fast rates of permeation and are restricted to solutes where *P_m_* < 10^−2^ cm/s (26).

These techniques have a range of capabilities and limitations. Simple planar-bilayer cells can be used across a range of solutes, provided that permeation is slower than the solution mixing time. Liposomes loaded with fluorescent reporters can also be used with slowly-permeating solutes that can be measured by fluorescence spectroscopy. Microelectrode methods can be used when the concentration of the solute can be measured electrochemically or can be measured indirectly by a change in an equilibrium (e.g., pH), although this can involve complicated kinetic models due to multi-species equilibria and unstirred layers at the membrane-water interface (27; 13; 28). Magnetic resonance methods, like EPR and NMR, can only be used for select solutes. The challenges associated with experimental permeability measurements have encouraged the development of molecular simulation methods that can support experimental results and provide insight into trends in permeability.

### 1.2. Permeability Models

Understanding the relationship between the chemical properties of the solute and its permeability is important for predicting the toxicity and pharmacokinetics of a compound (29; 30; 31). Models for predicting membrane permeability date back well over 100 years; publications by Meyer in 1899 and Overton in 1901 established the relationship between high rates of nonfacilitated membrane permeation and the hydrophobicity of the solute (32; 33). Walter and Gutknecht quantified this observation by showing that there is strong linear correlation between the log of the permeability coefficient and the log of water-alkane partition coefficients for a wide range of neutral solutes (34). This supports the Meyer–Overton rule that the rate of permeation is proportional to the relative solubility of the solute in the apolar membrane interior vs an aqueous solution.

Several experimental and theoretical models have been developed to predict permeability (35; 36; 37; 29; 30), with varying degrees of success, but the advent of computer simulations have led to the most significant developments. Molecular simulation methods for lipid bilayers have made it possible to study the permeation of solutes without direct empirical inputs. These simulations have provided atomic-scale interpretations of these data and quantitative interpretations of permeability trends in terms of the solution thermodynamics and dynamics. Empirically-based principles of permeation like the Meyer–Overton rule can now be interpreted within a rigorous physical framework. This review presents some of the research on the simulation of nonfacilitated permeation over the last 20 years. Interested readers may also be interested in a recent review by MacCallum and Tieleman on simulations of small molecules interacting with lipid bilayers (38) and a review by Orsi and Essex on modeling permeability (39).

## 2. Development of the solubility-diffusion model

To develop a quantitative theory for nonfacilitated permeation, the membrane is described as a fluid environment that the permeant passes through by Brownian motion. This model is consistent with direct molecular dynamics simulations of membrane permeation. Fig. 1 shows the trajectory of a O_2_ molecule permeating through a DPPC bilayer. The solute undergoes an effectively random walk along the *z*-axis before exiting the opposite side of the membrane. The solute also undergoes significant lateral motion inside the bilayer in the *xy* plane, but a rate theory can be developed based on the net flux (*J*) of the solute along the *z*-axis alone.

**Figure 1:**
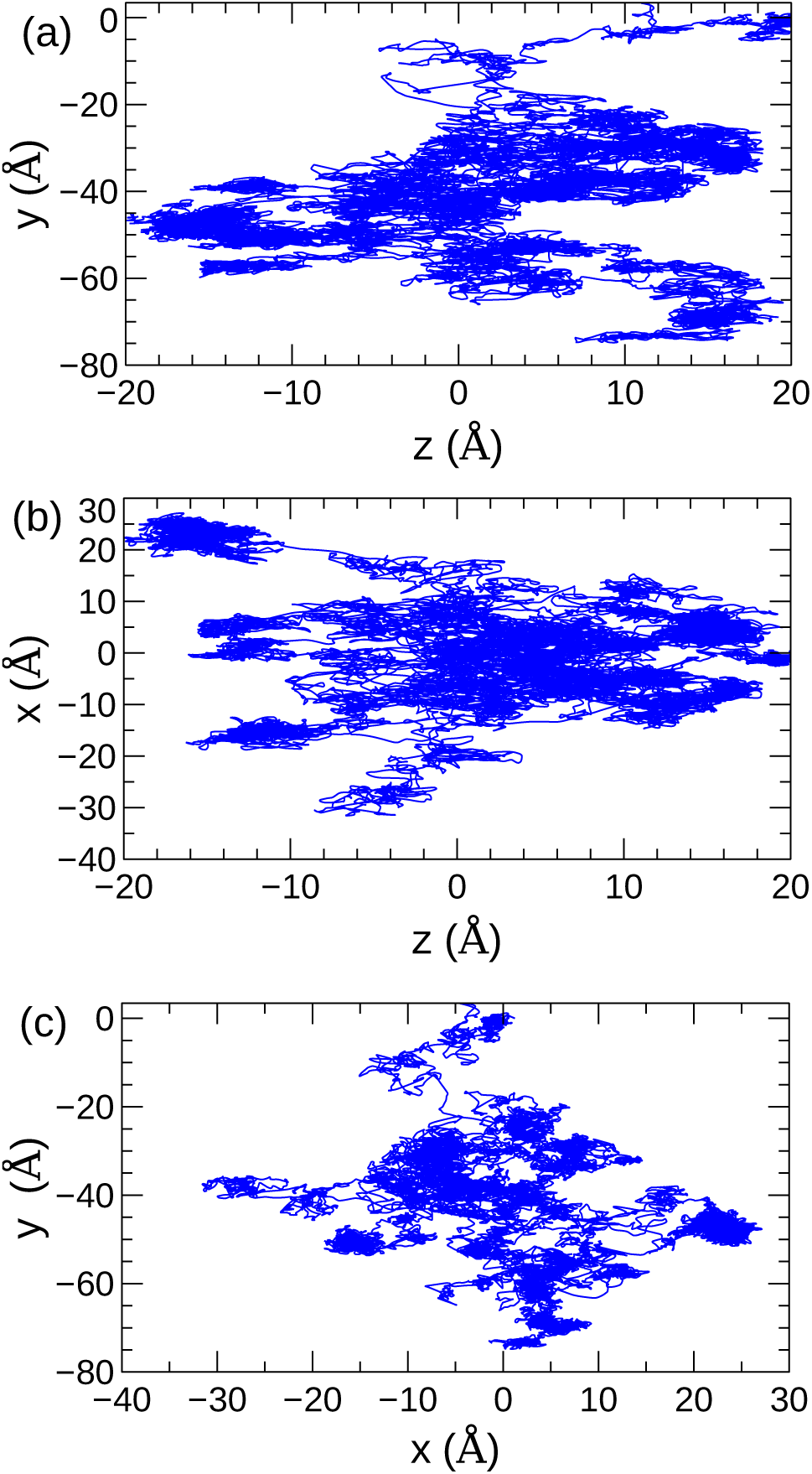
2D representations of a 300 ps trajectory of an O_2_ molecule permeating from the lower interface (*z* = −20 Å) to the upper interface (*z* = 20 Å) of a DPPC bilayer. The *z* axis corresponds to permeation across the bilayer, while the *x* and *y* axes are in the plane of the bilayer–water interface. The trajectory plotted in (a) and (b) show permeation consistent with Brownian motion, with the solute moving back and forth along the *z* axis before reaching the opposite interface. Plot (c) shows that there is significant lateral diffusion of the solute in the *xy*-plane during permeation.

The dynamics of the solute are complicated by the inhomogeneity of the bilayer, which varies in chemical composition and density along the transmembrane axis, *z*. In the inhomogenous solubility-diffusion model model, both the solute diffusivity *(D(z))* and potential of mean force (PMF, *w(z))* vary as a function of the position of the solute along the *z* axis. *w(z)* is related to the solubility of the solute at *z*, *K*(*z*) = exp(−*w(z)/k_B_T*), so this model is commonly referred to as the solubility-diffusion model.

The one-dimensional Nernst–Planck equation for a neutral solute in an inhomogeneous medium (40; 41) provides an expression for the flux through a unit area of the membrane at depth *z* (J(*z*)),

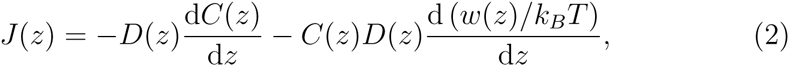

where *C*(*z*) is the concentration of the solute.

Multiplying both sides of the equation by 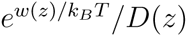 gives,

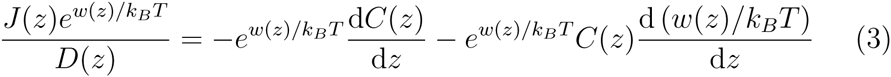

The right hand side of this equation can be combined into a single differential,

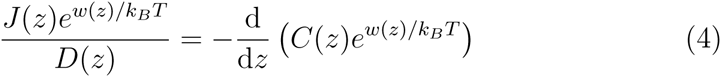

To determine the flux across the entire bilayer, this equation is integrated with respect to *z* over the interval [−*L/2,L/2]* where *L* is the width of a bilayer centered at *z* = 0,

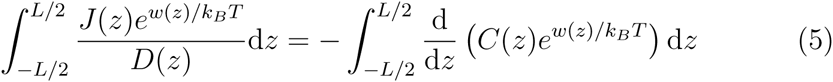

Under steady-state conditions, the flux at any value of *z* will be constant, so *J* can be factored out of the integral. Through rearrangement and simplification, the equation becomes,

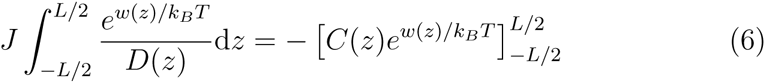

Isolating *J* on the left hand side and evaluating the bounds on the integral on the right hand side gives,

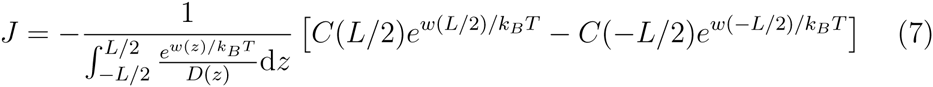

By definition, *w(z)* is zero in the solution outside the bilayer, so *w(L/2)* = *w(−L/2)* = 0. This gives the final form of the equation for the flux,

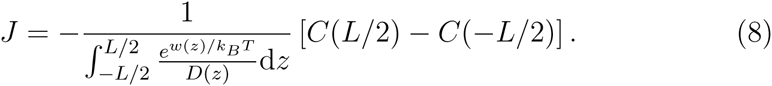

C(−L/2) − C(L/2) is the concentration difference across the bilayer in the direction of the positive flux, Δ*C*. By comparison to Eqn. 1, the permeability coefficient is,

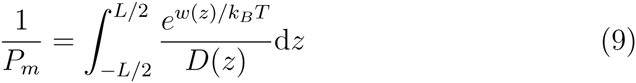

Alternative derivations are available in Diamond and Katz (42) and Marrink and Berendsen (43; 44). An elaboration that includes the orientational degrees of freedom of the solute is presented in Ref. 45.

Several assumptions are made in this derivation. The flux is assumed to be under steady state conditions with a negligible concentration gradient across the membrane. The membrane must be sufficiently laterally homogeneous so that meaningful averages of *w*(*z*) and *D(z)* can be determined. For example, a permeability rate that is governed by the formation of a water-pore that allows passage of the solute would not be well described by this model (7; 46). The solute dynamics are described by Brownian motion, which is only valid for individual solute molecules moving through high-friction environments on a reasonably flat free energy surface. The permeation of a solute with a variable number of associated water molecules would also be affected by the rates of hydration/dehydration inside the membrane and is not entirely consistent with this model.

To calculate *P_m_* using this theory, *D(z)* and *w*(*z*) must be determined for the full width of the membrane. These data can then be used to evaluate the integral in Eqn. 9 numerically. The bounds of the integration are also ambiguous because the membrane interface is diffuse; *w*(*z*) and *D(z*) must be calculated over a broad range so that the bounds on the integral can be determined based on when these values plateau to their bulk solution values. The practical aspects of calculating these profiles are presented in the following sections.

## 3. Practical aspects

### 3.1. Simulation systems

Simulations of membrane permeability are typically performed with periodic unit cells. A planar bilayer is constructed in the cell to form a lamellar system. Water layers are added above and below the bilayer. By convention, the bilayer interface extends along the *xy* plane and the *z*-axis corresponds to the transmembrane axis (a.k.a., the bilayer normal). Recent simulations commonly include 60–200 lipids (47; 48; 49). The water layers must be sufficiently thick so that periodic membrane–membrane interactions are minimized and so *D*(*z*) and *w*(*z*) can be calculated at distances that are sufficiently far from the water-lipid interface to establish a reference (i.e., *w*(*z*_bulk_) = 0). Studies have used simulation cells with various sizes, but bilayer surface areas in the (40 − 60)^2^ Å^2^ range and cell heights of 80 − 110 Å in the *z* dimension are common. A typical simulation cell is shown in Fig. 2.

**Figure 2:**
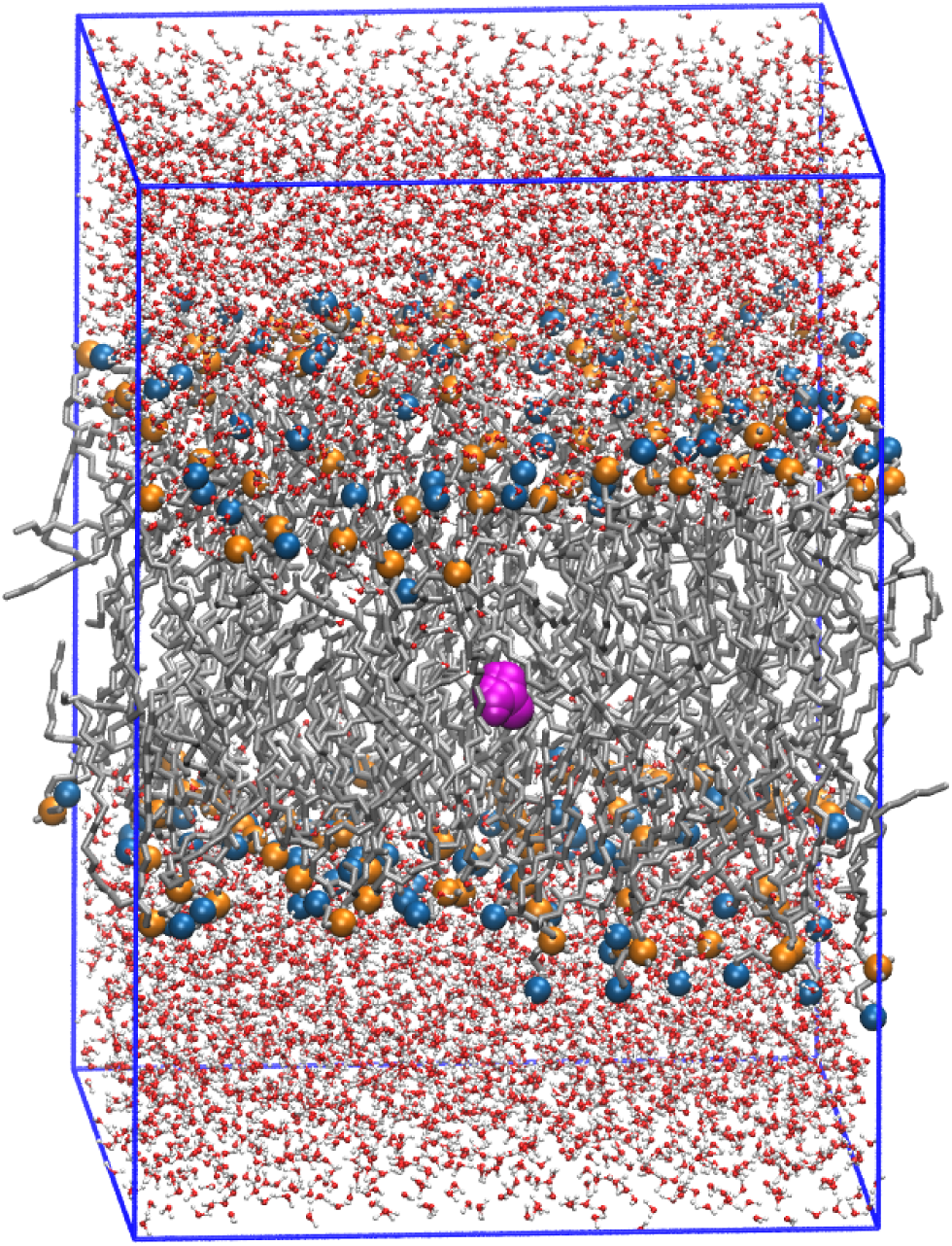
An example simulation cell used to simulate membrane permeation. A bilayer section with a 62 Å × 62 Å surface area and containing 128 1,2-dimyristoyl-sn-glycero-3-phosphocholine (DMPC) lipids is oriented in the *xy* plane of the cell. Lipid tails are rendered in gray. Lipid heads are rendered in blue (choline) and orange (phosphate). The bilayer is surrounded by water layers (red and white). A permeating molecule of urea is at the center of the bilayer (magenta).

A wide range of force fields have been used in recent permeation simulations. The TIP3P (50) and TIP4P (51) water models are both popular. The Berger (52) and CHARMM36 (53) lipid models have both been used successfully. The permeating solutes have been described using specific force fields or generalized force fields like GAFF (54), OPLS (55), or CGenFF (56). Polarizable and coarse grain models are also available. These are discussed in Sections 5.1 and 5.2.

### 3.2. Calculation of the Potential of Mean Force

The potential of mean force of the permeation of the solute along the *z*-axis (*w*(*z*)) can be calculated using a range of free energy methods. The first PMFs calculated by Marrink and Berendsen used (43),

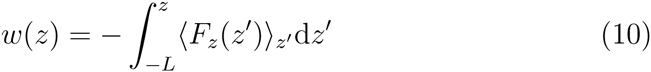
where *F_z_*(*z′*) is the force on the solute along the *z*-axis when the solute is constrained at *z*′.

To sample the ensemble average of *F_z_*(*z′*), (〈*F_z_*〉(*z′*), the equations of motion must be modified so that the solute is constrained to a value of *z′*. The *z*-component of the force on the solute by the environment are averaged over long MD simulations at a series of positions spanning the membrane. The integral is numerically integrated to yield *w*(*z*).

Most recent permeability simulations use umbrella sampling to calculate *w*(*z*) (57; 58). In this method, a series of simulations are performed with the solute restrained relative to the center of mass of the membrane using a harmonic potential,

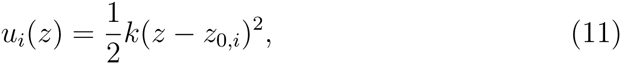
where *u_i_*(*z*) is the potential energy of the restraint for simulation *i, k* is the force constant of the restraint, and *z*_0_,*_i_* is the reference position along the *z* axis the simulation is biased to sample near.

These simulations are performed with values of z_0_,*_i_* separated by regular intervals, such that a solute is simulated extensively at every point spanning the membrane. The optimal choices of window spacing and force constant (*k*) depends on the membrane composition, temperature, and solute. Recent studies have used window spacings that vary between 0.2 and 1.0 A separations and force constants in the *k* = 1.2 − 7.2 kJ mol^−1^ A^−2^ range (59; 48; 47). In systems where the PMF varies sharply with *z*, sections of the PMF may require more extensive sampling (60), so additional windows at smaller spacings and stronger force constants can be added in these regions (59). *G*(*z*) can be constructed from the distributions sampled in these simulations using techniques like Weighted Histogram Analysis Method (WHAM) (61; 62; 63) or the multistate Bennett acceptance ratio estimator (MBAR) (63).

Neale et al. showed that umbrella sampling simulations for the partitioning of arginine and isoleucine side chain models into a bilayer required very extensive simulations to achieve sub-kcal convergence (60). This was particularly serious for the charged arginine group, which required a 125 ns equilibration and *μ*s length sampling per umbrella sampling window to eliminate systematic errors. This was attributed to distortions in the bilayer that occur over long timescales. Even the calculation of permeation PMFs of small nonpolar solutes requires equilibration simulations that are several nanoseconds long and a sampling period greater than 10 ns. Performing several independent simulations from different initial configurations can help identify if the PMF has reached convergence.

The slow convergence of permeation umbrella sampling simulations can be addressed by enhanced sampling techniques. Hamiltonian-exchange molecular dynamics allows neighboring windows of an umbrella sampling simulation to periodically exchange coordinates (64; 58). This can be applied to the calculation of membrane permeation PMFs by allowing exchanges of neighboring umbrella sampling windows along the *z* axis. For example, exchanges are periodically attempted between the simulation where z_0_ = 4 Å and the simulation where z_0_ = 5 Å. Neale et al. (47) developed a Hamiltonian exchange method for membrane permeation PMFs based on the virtual replica exchange method of Rauscher et al. (65). This method was found to have 300% better sampling efficiency than conventional umbrella sampling simulations.

Huang and Garcia applied the Replica Exchange with Solute Tempering (REST) enhanced sampling method to the simulation of lipid bilayers (66). In this technique, the simulation of the bilayer is allowed to undergo exchanges with simulations where the solute and bilayer are coupled to different thermostats and lipid-lipid interactions are biased. This allows rapid lateral diffusion of the lipids and could improve the sampling of the configurational space of the permeating solutes. Jaabeck and Lyubartsev have found that metadynamics can help sample solute conformational degrees of freedom, which allows better sampling of the PMF of permeation (67).

The permeation of large amphiphilic solutes is complicated by the dynamics of secondary degrees of freedom. Even for small amphiphiles, like alcohols, the solute must undergo a “flip-flop” transition, where the hydrophilic head reorients to be directed towards the opposing membrane interface when it passes through the bilayer interior. The transitions between these orientations can require time scales greater than nanoseconds, so a conventional umbrella sampling simulation along the *z* axis may not provide an accurate description of the permeation.

This sampling issue with amphiphilic solutes can be addressed by biasing the simulation to sample the orientational degrees of freedom of the solute. Jo et al. calculated the PMF for the transition of cholesterol from one bilayer leaflet to the other by a 2D umbrella sampling simulation of position of center of mass along the *z*-axis and the tilt angle of the solute with respect to the plane of the bilayer (68). Comer et al. developed a method that incorporates the dynamics of the solute along *z* and its orientation with respect to the *xy* plane of the membrane (69). Similarly, Parisio et al. developed an extension to the solubility-diffusion model where the rotation of the solute is explicitly included (45). Based on this model, Pariso, Sperotto, and Ferrarini concluded that the predominant pathway involved reorientation of the steroid while in the top leaflet so that its headgroup points towards the opposite side of the bilayer, then permeation along the *z* axis to enter the lower leaflet (70).

### 3.3. Calculation of the diffusivity profile

The solubility-diffusion model requires the calculation of the diffusion coefficient of the solute as a function of the position of the solute along the *z*-axis (*D*(*z*)). The Einstein equation allows the diffusion of the solute to be calculated by performing an MD simulation and calculating the mean square displacement of the solute along the *z*-axis (71),

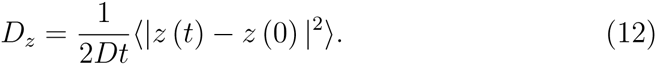

Alternatively, the Green–Kubo relation allows diffusion coefficients to be calculated from the integral of the velocity autocorrelation function (72),

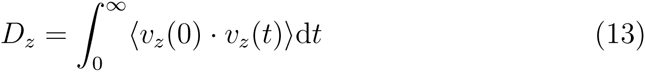

The inhomogeneity of a lipid bilayer makes it difficult to apply these methods to the calculation of *D*(*z*). These methods are not appropriate for calculating *D*(*z*) because they assume the solution is homogeneous, so they cannot capture the variation of *D* with the position of the solute along *z*.

Hummer developed a method to calculate position-dependent diffusion coefficients by sampling the frequency of transitions between positions along a coordinate in an equilibrium MD simulation (73). The variation of solubility of the solute in the bilayer is a challenge for this method. If the solubility of the solute is low at any point of the profile (i.e., *w*(*z*) is large), transitions across this part of the profile in an equilibrium MD simulation will be rare. Very extensive MD simulations would be needed to directly sample diffusion across the full bilayer except for the most permeable solutes.

One direct method of calculating *D*(*z*) uses a relation from fluctuation–dissipation theory to calculate *D*(*z*) using the autocorrelation function (ACF) of the force on the solute when it is constrained at position *z* along the axis (72),

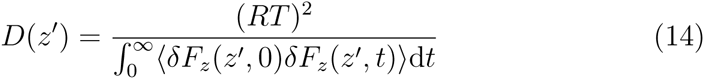

Here, *δF_z_*(*z*′, *t*) is the deviation of the force on the solute along the *z* axis at position *z*′ from the average force (*δF_z_*(*z, t*) = *F_z_*(*z, t*) − 〈*F_z_*(*z*′)〉).

In practice, these calculations make use of the same type of simulations that can be used to calculate the PMF from the integral of the average force (Eqn. 10). The drawback of this method is that most MD codes would have to be modified in order to do these calculations. The equations of motion of the MD integration algorithm must be modified to constrain the solute to a particular value of *z* (74). Most standard simulation codes must also be modified to output the time series of the force on a single molecule.

Alternative techniques for calculating D(*z*) are provided by solutions to the Generalized Langevin Equation (GLE) for a harmonic oscillator. Consider a simulation where the permeant is restrained at some value of *z* along the transmembrane axis using a harmonic potential (i.e., Eqn. 11). The motion of the solute can be modeled as a harmonic oscillator where the rest of the system serves at its frictional bath. The GLE expression for the dynamics of this oscillator is,

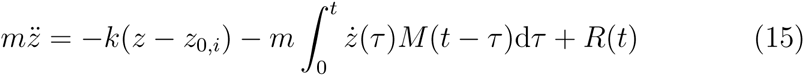
where *R*(*t*) is the random force that originates from the degrees of freedom orthogonal to *z*. *M*(*t*) is the memory kernel, which reflects the frictional forces on the solute. Beginning from this equation, Woolf and Roux derived an expression for the diffusion coefficient at 〈z〉*_i_* in terms of the Laplace transform of the velocity 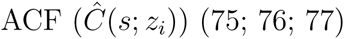 (75; 76; 77).

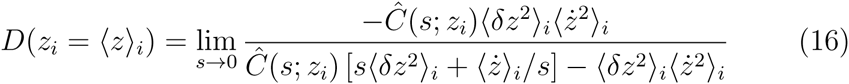

Because this equation has a singularity at *s* = 0, the limit cannot be taken directly. Instead, the Laplace transform must be performed numerically for a series of values of s and then the value of *s* = 0 must be extrapolated from these points.

Hummer derived a simpler expression based on this theory that relates *D*(*z*) to the variance of *z* and the correlation time (*τ*) (73),

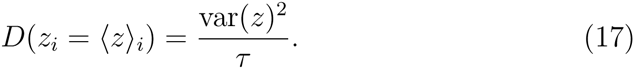
τ is calculated from the integral of the position ACF (*C_zz_* (*t*)).

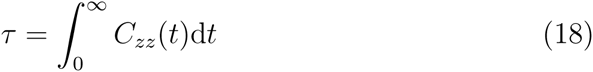

*C_zz_* can be calculated by direct summation of pairs over the time series (78),

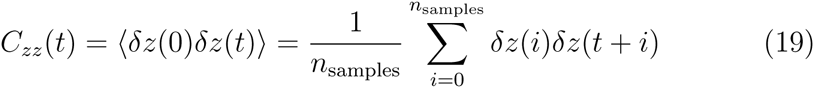
where *δz*(*t*) = *z*(*t*) − 〈*z〉.*

Zhu and Hummer presented a further simplification that avoids calculation of a correlation function (79). In this method, τ is calculated from the variance of the mean value 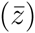 and the interval of the time series data points (Δ*t*),

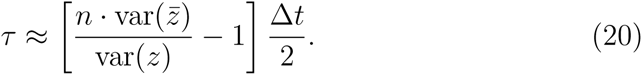

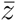 can be calculated using the block averaging method of Flyvbjerg and Petersen (80). This provides perhaps the simplest method for calculating *D*(*z*) from time series data, although the analysis of the time series to calculate *D*(*z*) using any of these methods is small in comparison to the cost of performing the MD simulations to generate the time series.

A key advantage of these methods is that they only require that the solute be restrained at a point along *z* with a harmonic potential with a reference position relative to the center of mass of the bilayer. This type of restraint is simple to implement and is available in most molecular dynamics codes. The diffusivity profile of a solute can be calculated by performing a series of MD simulations where the solute is restrained to positions at regular intervals along the *z* axis (e.g., 1 Å apart). This procedure is directly analogous to the umbrella sampling simulations using harmonic restraints that are used to calculate *w*(*z*). In principle, both properties could be calculated from the same simulations. A time-series of the *z* position of this solute is collected from each simulation, which is then used to calculate the ACF.

In practice, the calculation of *D*(*z*) is highly sensitive to the equilibration of the system and the length of sampling. Multiple simulations that are several nanoseconds long are needed to calculate well-converged ACFs. The spring constants of the harmonic potential (*k*) can affect the results because the description of the system with the GLE is based on the system acting as a harmonic oscillator in a frictional bath. Deviations away from this harmonic-oscillator model due to the effect of a rough underlying free energy surface can introduce biases. Hummer also showed that *D*(*z*) varies with the length of the interval the ACF is integrated over (73). Lastly, irregularities in the bilayer structure and the formation of water clusters around the solute can lead to large variations in *D*(*z*). Improved methods for calculating position-dependent diffusivity profiles are an active subject of investigation (81; 82).

## 4. The solubility-diffusion model in practice

### 4.1. Interpreting permeation profiles

#### 4.1.1. Bilayer regions

The diffusion and PMF profiles calculated by Marrink and Berendsen illustrated how the bilayer environment varies with depth (43; 83). The bilayer was divided into four overlapping regions that have different structural and chemical interactions with solutes, so the solutes will diffuse at different rates and have different solubility in these regions. The center of the bilayer is used as the point of reference (*z* = 0). The given ranges of *z* are approximate values for the DPPC bilayer example used in this review.

- Region I: |*z*| > 25 Å. The bulk solution. This region is primarily liquid water, with a small component of lipid headgroups.
- Region II: 17 Å < |*z*| < 25 Å. The lipid headgroups. These are typically solvated by water molecules.
- Region III: 10 Å < |*z*| < 17 Å. The upper portions of the lipid tails, acyl groups, and lower portions of the head groups.
- Region IV: |*z*| < 10 A. The interface of the lipid tails of the two opposing monolayers at the center of the bilayer

These regions will be used to describe sections of the potential of mean force and diffusivity profiles discussed in the following sections. Marrink and Berendsen compared the PMFs of O_2_ and H_2_O as the prototypical examples of hydrophobic and hydrophilic permeants, respectively (83). These calculations were reproduced here using current models and simulation lengths (Fig. 3). The technical details of these simulations are included in Appendix A.

**Figure 3:**
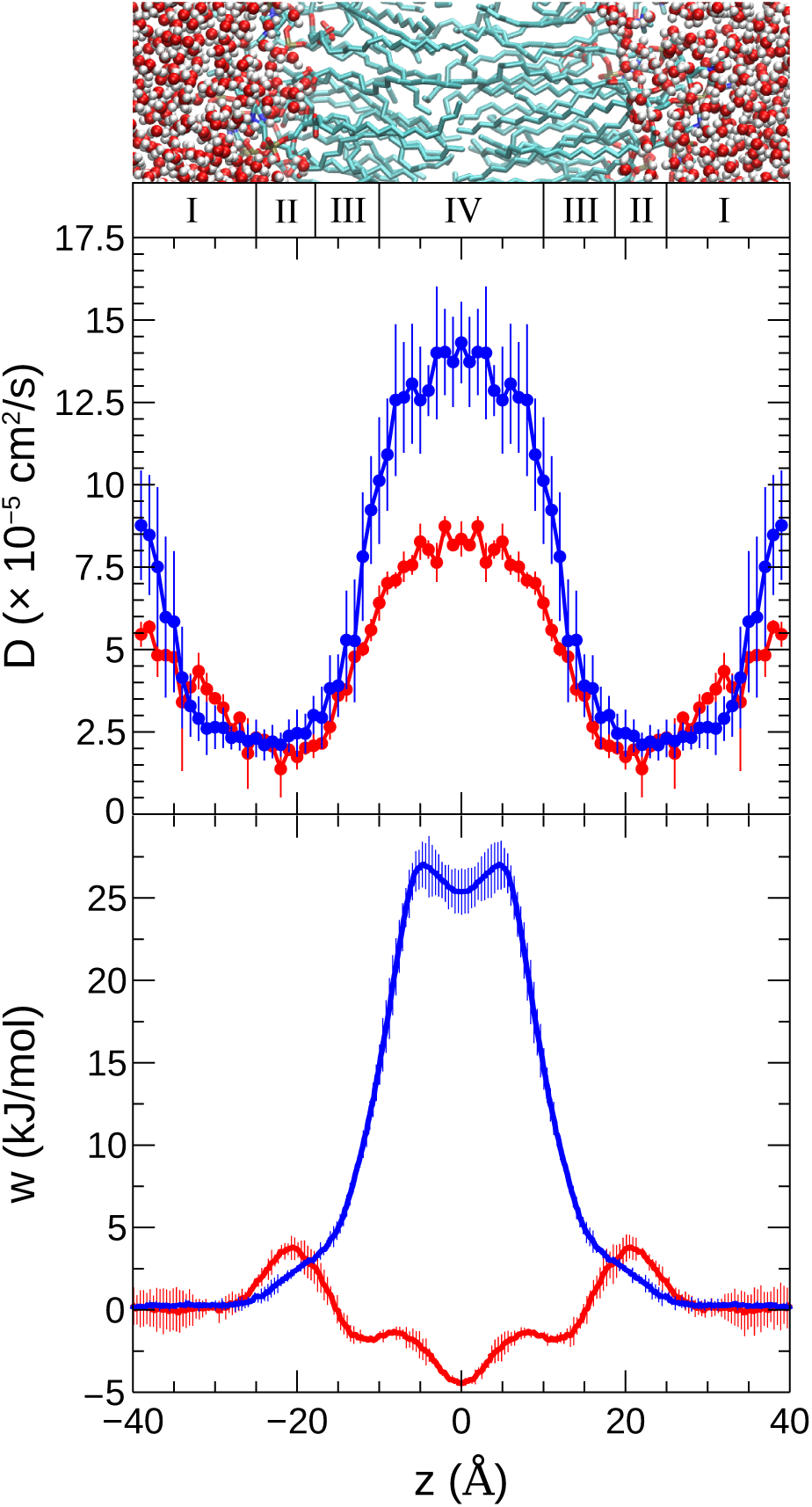
The diffusivity (*D*(*z*), top) and potential of mean force (*w*(*z*), bottom) of H_2_O and O_2_ permeating through a DPPC bilayer. The curves for H_2_O are shown in blue while the curves for O_2_ are shown in red. Simulations were performed at 323 K. The CHARMM36 lipid model and TIP3P water models were used. The PMF profiles were calculated from 20 ns H-REMD umbrella sampling simulations. The diffusivity profiles were calculated from the average of six 4 ns MD simulations using Eqn. 17.

#### 4.1.2. Potential of Mean Force

The PMF, *w*(*z*) (a.k.a., Δ*G*(*z*)), is the reversible work needed to move the solute from solution to the position *z* on the transmembrane axis. The PMF corresponds to the relative solubility of the permeant in solution vs the membrane interior; a positive PMF indicates that the solute is less soluble at this point in the membrane than it is in bulk solution, while a negative PMF indicates that it is more soluble. A highly positive PMF inside the membrane results in a small permeability coefficient, consistent with a low probability of the solute partitioning inside the bilayer to this position.

Hydrophobic solutes like O_2_ have a small increase in the PMF at the water-lipid interface. The PMF drops once the solute reaches the ester/tail regions (Regions III & IV). For many hydrophobic solutes, the PMF reaches a minimum at *z* = 0 Å, indicating that these hydrophobic compounds will concentrate at the center of the bilayer. The thermodynamic driving force for this effect was investigated by MacCallum et al. by decomposing the PMF of hexane permeation into enthalpy and entropy components (59). Although the PMF across the bilayer was broad and flat, with a minimum at the center, the entropy and enthalpy components of the PMF varied significantly as the solute moved across the bilayer. The enthalpy component disfavors partitioning of hexane at the center of the bilayer where it has weaker intermolecular interactions with the lower-density lipid tails, but this is counteracted by a more favorable entropy term because the solute has greater configurational freedom in the fluid-like bilayer center.

The PMF of water is typical for the permeation of a polar solute. The PMF of permeation for hydrophilic solutes do not show large changes in the headgroup or ester regions. These regions are easily accessible by the solvent, so the solutes are largely hydrated in these regions and can have stabilizing interactions with the head groups. The PMF increases abruptly when the solute enters the lipid chain region of the membrane (Region III). The PMF reaches a maximum in the lipid tail region (Region IV) that extends over 10Å.

Permeation of charged and hydrogen-bonding solutes shows the most distinct PMFs, with a sharp peak at the center of the bilayer. These compounds remain partially hydrated by a finger of water molecules that extends from the headgroups into Regions III & IV. In permeation simulations of ionic compounds like Na^+^, Cl^−^ (84), methylammonium (85), and methylguanidinium (86; 60; 87), the solute remains hydrated by a cluster of water molecules even at the center of the bilayer. Calculating the PMF is particularly challenging because various hydration states must be sampled. Because of this variable hydration, the motion of the solute cannot be reliably described as Brownian motion of a single particle, so the solubility-diffusion model is not necessarily a valid means to calculate a permeability coefficient. Nevertheless, the PMF can still be interpretively useful.

In one of the most extensive studies of nonfacilitated ion permeation, Vorobyov et al. used simulation and electrophysiological experiments to explore the permeation mechanisms of Na^+^, K^+^, Cl^−^, and GuanH^+^ (88). The PMFs were largely similar despite differences in size and charge, yielding similar membrane permeabilities that agree semiquantitatively with experiments in terms of magnitude and selectivity. The hydration of the ions inside the membrane negates the differences in the ion solvation energy that would be dominant in a traditional solubility-diffusion permeation mechanism. This led them to propose that permeation of ionic compounds occurs through a distinct, ion-induced permeation mechanism, where the ion induces a deformation in the membrane that allows the ion to permeate without being desolvated.

### 4.2. Connection to partition coefficients and the Meyer-Overton rule

A central tenet of the solubility-diffusion model is that the concentration at a given point, *z*, is determined by the equilibrium between the solute partitioning to the position *z* in the bilayer layer and the solution.

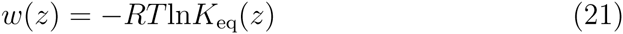

Due to the exponential dependence on *w*(*z*) and the tendency for *w*(*z*) to have a broad plateau at the center of the membrane, the permeability coefficient of a solute is largely determined by the value where *w*(*z*) is a maximum. This suggests a proportionality between the log of the permeability coefficient and the equilibrium constant for the solute,

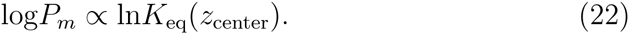

As a solvation environment, the interior of the membrane is similar to a liquid alkane, so the equilibrium constant between the solvent and the center of the membrane will in turn be proportional to the alkane/water partition coefficient (*P*),

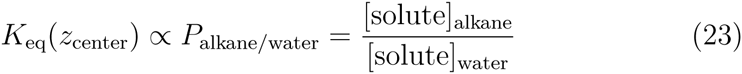

Note the difference between the permeability coefficient, *P_m_*, and the partition coefficient, *P*. *P_m_* gives the rate of flux of a solute across the membrane through Eqn 1 and has units of distance per unit time. The partition coefficient is a measure of the relative concentration of the solute in two phases and is dimensionless.

The octanol/water and hexane/water partition coefficients are commonly used as measures for the degree to which a solute will partition into the membrane interior. It has been suggested that the octanol/water partition coefficient is a better measure for partitioning into the lipid headgroups than the membrane interior (89). Walter and Gutknecht showed that the hexadecane/water partition coefficient has the strongest correlation to membrane permeability, with a correlation coefficient of *r* = 0.950 (34). Based on this, the permeability coefficient and the alkane/water partition coefficients are closely related (i.e., log*P_m_* ∝ log*P*_hexadecane/water_), which provides a theoretical basis for the Meyer–Overton rule.

#### 4.2.1. Diffusivity profiles

The diffusivity profiles of solutes follow some general trends. The diffusivity is high at the center of the membrane where the lipid tails are disordered and fluid-like. The diffusivity drops significantly in Region III, where the lipid tails are ordered and undergo slow rearrangements (90). The diffusivity remains at a depressed value in the headgroup/water interface region, but increases again when the solute enters the solution (Region I). It should be noted that the diffusivity in the solution region will generally be overestimated by a factor of 2–3 when the TIP3P water model is used because this model has an erroneously low viscosity (91).

The diffusivity of larger permeants tends to be smaller than for smaller permeants. In Fig. 3, the diffusivity of the diatomic O_2_ is systematically smaller than for H_2_O, which has only one non-hydrogen atom. This can be interpreted using the Stokes-Einstein equation (71),

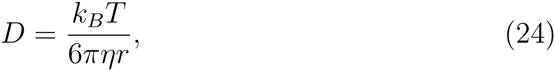
which predicts that the diffusion coefficient of a solute in solution is inversely proportional to the molecular radius, *r*. The Stokes–Einstein diffusion coefficient also has an inverse dependence on the viscosity of the solvent (*η*), which predicts that solutes would diffuse more rapidly in low-friction regions of the bilayer.

Broadly speaking, the diffusivity of a solute varies by a factor of 5–6 across the bilayer. While this is a significant difference, the solubility-diffusion model permeability coefficient is only linearly dependent on the *D*(*z*). As a result, the permeability of a solute is less sensitive to *D*(*z*) than it is to *w*(*z*), which has an exponential dependence. Furthermore, the high diffusivity in Region IV largely cancels the low diffusivity in Regions II & III. This generally supports the strategy of many researchers who have interpreted trends in permeability based only on the PMF; however, quantitative comparison to experimental permeability coefficients still require accurate calculations of the diffusivity profiles.

### 4.3. Applications of the solubility-diffusion model

The 1994 paper by Marrink and Berendsen on the calculation of permeability coefficients from molecular simulation has been cited more than 500 times as of 2015 (43). The model has been used to describe the permeation of solutes ranging from gases to nanoparticles. Table A.2 in Appendix A summarizes 85 papers where membrane permeability is studied using molecular simulation.

Although many studies have focused on qualitative analysis of membrane permeability, there are many examples of quantitative calculations using the solubility-diffusion model. Orsi and Essex reported one successful example where the permeability of *β*-blocker drugs was calculated (92). A dual-resolution model was used to calculate permeability rankings of alprenolol, atenolol, pindolol, progesterone, and testosterone through DMPC lipid bilayers. In general, the permeability coefficients were predicted within an order of magnitude.

Another study by Riahi and Rowley employed the solubility-diffusion model to explain the high membrane permeability of hydrogen sulfide (H_2_S). In this study, the authors employed all-atom polarizable molecular dynamics simulations to compute the permeability of H_2_O and H_2_S through a DPPC lipid bilayer. The calculated membrane permeability coefficients for both permeants were in good agreement with the experimental values.

Holland and coworkers studied the permeability of molecular oxygen and trace amine p-tyramine (charged and neutral) through model phosphatidylcholine (PC) lipid bilayers (93). The permeability of oxygen was studied at two temperatures, 323 K and 350 K, while that of p-tyramine was studied at 310 K. The PMF and diffusion coefficient, *D*(*z*), for the permeants were obtained using non-equilibrium work methods, which in turn were used to calculate the permeability coefficient of the permeants using the solubility-diffusion model. The calculated permeability coefficients for the selected permeants were in good agreement with experimental permeability of similar lipid bilayers.

### 4.4. Effect of cholesterol content

One of the most significant contributions of the solubility-diffusion model has been in understanding the effect of cholesterol on membrane permeability. Cholesterol is a major component of eukaryotic cell membranes and has a large effect on membrane permeability. Generally, solutes permeate membranes with higher cholesterol content at lower rates (94; 95; 96).

One of the earliest simulation studies on the effect of cholesterol content on membrane permeability was reported by Jedlovszky and Mezei (97). Simulations of 8 solutes (H_2_O, O_2_, CO, CO_2_, NO, NH_3_, CHCl_3_, and formamide) permeating through DMPC bilayers with varying cholesterol concentration pointed to changes in the PMF as the cause of the reduced permeability. A later study by Hub et al. found that the PMF for the permeation of ammonia and carbon dioxide was increased when the cholesterol content of the membrane was high (98). The effect of cholesterol on the permeation of drug molecules has also been studied. Eriksson and Eriksson investigated the influence of cholesterol on the membrane permeability properties of the photodynamic drug, hypericin, and its mono- and tetra-brominated derivatives. The authors found that the calculated rate of permeation was lower at high cholesterol content due to higher PMF barriers (99).

Several simulation studies have examined the effect of bilayer cholesterol content on water permeation, which also decreases at high cholesterol content (23). Saito and Shinoda explored the effect of cholesterol content on the water-permeation PMFs of DPPC and PSM lipid bilayers (100). They found that with increasing cholesterol concentration, both DPPC and PSM lipid membranes displayed an increased PMF for water permeability; although the effect was more prominent in the PSM bilayers with 30 mol % of cholesterol content. Similarly, Issack and Peslherbe examined the influence of cholesterol on the PMF and diffusivity profiles for transmembrane water permeation (101). They found that the PMF in Region III was significantly increased when the cholesterol content was high. A small decrease in diffusivity in Region IV had a secondary contribution.

Wennberg et al. reported a comprehensive study of the effect of cholesterol on permeation (48). The PMFs for solute permeation across 20 different lipid membranes containing one of four types of phospholipids (i.e., DMPC, DPPC, POPC, POPE) and cholesterol content that varied from 0 to 50 mol % were computed. The general trend from this study was that the PMFs for solute permeation were increased when the cholesterol content was high, particularly in Region III, where the ring structure of cholesterol preferentially partitioned. Cholesterol is relatively rigid and packs tightly into the lipid tails, resulting in strong London dispersion interactions between the lipids and the cholesterol. A permeating solute disrupts these interactions and is generally unable to pack into the lipid tails as tightly as can cholesterol. As a result, the PMF for a solute permeating through Region III of a cholesterol-containing bilayer is increased, which decreases the rate of permeation. A subsequent study by Zocher et al. found the diffusivity of permeating solutes were generally independent of cholesterol content and changes in the PMF were the dominant effect on permeation rates (102).

#### 4.4.1. Lipid composition

The effect of lipid composition of permeation has also been studied computationally. These studies have primarily used model amino acid side chains as the solutes because the partitioning of side chains into the interior of a membrane is relevant to membrane protein structure and function.

One notable study by Li et al. investigated the role of membrane thickness in saturated phosphatidylcholine (PC) lipid bilayers (85). The authors calculated the PMFs for the permeation of arginine side chain analog MguanH^+^ across PC lipid bilayers with increasing thickness (DSPC, DPPC, DMPC, DLPC, DDPC). The authors proposed that an ion-induced defect mechanism is the principal means of arginine translocation within lipid bilayers with lipid tails up to 18 carbons in length (e.g., DSPC). For thicker bilayers, the solubility-diffusion mechanism is preferred.

The results for protonated methylguanidinum (MguanH^+^), a model for an arginine side chain, were particularly significant. For all lipid bilayers used in the study, similar membrane deformations were observed for MguanH^+^ translocation; where water and lipid head groups are drawn into the hydrophobic core of the bilayer. The free energy barrier for MguanH^+^ translocation in lipid bilayers display a systematic dependence on membrane thickness with predictable shifts between bilayers of increasing thickness. Narrow transmembrane pores and corresponding plateaus in free energy profiles are observed for MguanH^+^ translocation across thin membranes (DDPC, DLPC) suggesting a favorable energetic cost for burying an arginine residue into a thin bilayer; a finding that is consistent with experimental measurements (103). This work indicates that membrane thickness plays an important role in the membrane partitioning of charged amino acid side chains.

The effect of lipid headgroup and tail length of side chain partitioning was also studied by Johannson and Lindahl using molecular simulation (104). The PMFs for permeation of 8 side chain models (Asp, Glu, Arg, Lys, Ser, Trp, Leu, and Phe) was calculated for six different types of bilayer (DMPC, DOPC, DOTAP, POPC, POPE, and POPG) with different head and tail groups. The PMF barriers of polar and charged groups were higher and broader for thicker membranes, but PMF barriers for hydrophobic groups were lower. The charge of the lipid head group had a large effect on the permeation of charged compounds. The barriers for Glu and Arg side chains increased as the lipids were changed from cationic, to zwitterionic, to anionic headgroups.

## 5. Innovations in modeling permeation

### 5.1. Polarizable models

Permeating solutes pass from the polar aqueous phase through the nonpolar membrane interior. In the course of permeation, the dielectric constant changes from *ϵ* = 78 in bulk water to *ϵ* = 2 in the lipid tail region. Polarizable solutes can experience a large induced polarization when dissolved in liquid water. This polarization effectively disappears when the solute moves to the nonpolar membrane interior. For solutes that can hydrogen-bond with water, this change in polarization is even more significant because the solutes experience a large electric field from the water molecules. Most simulations to date use fixed atomic charges to describe electrostatic interactions, so they are incapable of describing the effects of induced polarization.

Force fields that include the effects of induced polarization are gradually becoming available. A variety of polarizable models for biomolecules have been developed, including the Drude (105; 106), CHarge EQuilibration Method (CHEQ) (107), and the Atomic Multipole Optimized Energetics for Biomolecular Simulation (AMOEBA) models (108). Drude models are attractive for bilayer simulations because of their efficiency; polarizability is incorporated by tethering charged “Drude” particles to non-hydrogen atoms and propagating their positions dynamically (see Fig. 4), so there is no additional iterative process to calculate the charge distribution.

**Figure 4:**
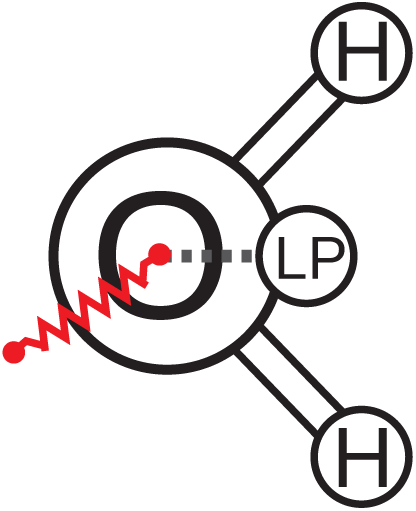
A schematic of the SWM4-NDP Drude polarize water molecule. Each water is comprised of three atomic centers (an oxygen and two hydrogens), a massless “lone-pair” particle is constrained to a position along the H–O–H bisector, and a negatively charged “Drude” particle that is tethered to the oxygen atom with a harmonic potential (red).

To date, only a handful of polarizable force fields for lipids are available. Davis et al. reported a CHEQ polarizable model for DPPC lipids (109). Separately, Robinson has developed a Drude model force field for cholesterol and sphingomyelin (110). Harder et al. developed a Drude polarizable model for DPPC to model the lipid monolayer-air interface (111). Chowdhary et al. refined this model to describe DPPC bilayers (112). It should be noted that this Drude DPPC model is actually less accurate in describing the lipid head group area and order parameters than the nonpolarizable CHARMM36 model (112), although this can likely be addressed by further refinement of the force field parameters.

A simulation of water permeation through a DMPC bilayer using a CHEQ model was reported by Bauer et al., who found the barrier in the PMF using various CHEQ polarizable models to be in the 19–23 kJ/mol range (113). This barrier is somewhat lower than the 26–30 kJ/mol barriers reported in other simulations (113). The permeability coefficient was not calculated in this study, so this result cannot be directly tested by comparison to the experimental value. There was a large change in polarization of the permeating water molecule; the dipole moment of water dropped from 2.6 D in bulk water to 1.88 D at the bilayer center.

Riahi and Rowley simulated the permeation of H_2_O and H_2_S through a DPPC bilayer using the Drude polarizable model (49). The average dipole moment of H_2_O decreased from 2.5 D to 1.9 D when it crossed into the center of the membrane interior, but the average dipole moment of H_2_S only decreased from 1.2 D to 1.0 D. This is contrary to the trend in solute polarizability, as H_2_S has a polarizability of 3.6 Å^3^, while water has a polarizability of 1.45 Å^3^ (Fig. 5). Counter-intuitively, molecules that are highly polarizable will not necessarily be strongly polarized in liquid water because high polarizability correlates to larger molecular radii. In the case of H_2_S, the modest polarity and large radius of the sulfur atom precludes hydrogen bonding or other kinds of close-range polarizing interactions with water. Induced polarization is most significant in molecules that have a large static dipole moment and form hydrogen bonds with water molecules.

**Figure 5:**
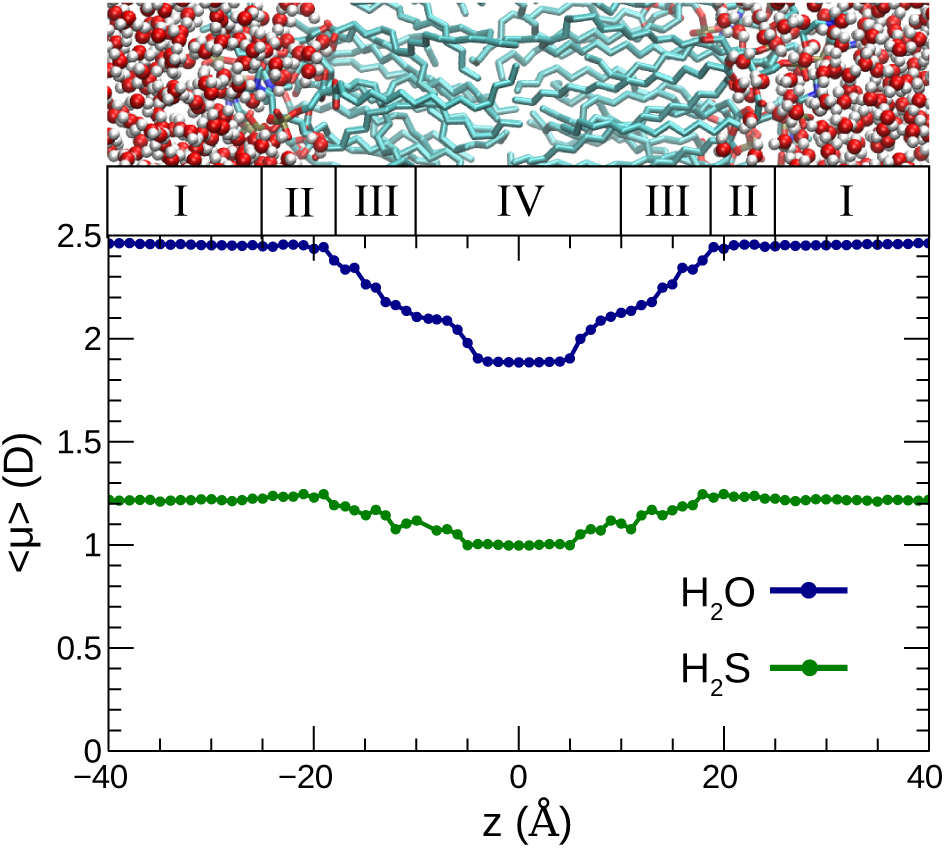
The dipole moments of H_2_O (blue) and H_2_S (green) solutes permeating across a DPPC bilayer. The average dipole of H_2_O is high in solution (〈*μ*〉 = 2.5 D) but drops to a value equal to its gas-phase dipole moment (*μ*_0_ = 1.8 D) at the center of the bilayer. H_2_ S experiences a much smaller change (〈*μ*〉 = 1.2 → 1.0 D) The values are averaged from the umbrella sampling windows, calculated with a Drude polarizable model. Data from Ref. 49.

Although the rigorous inclusion of induced polarization is desirable conceptually, the results from simulations using polarizable models are not necessarily more accurate. For instance, the water-permeation PMF calculated using the Drude model is very similar to that calculated using many non-polarizable models. Nonpolarizable force fields are often parameterized to reproduce the experimental solvation free energies of solutes. These energies correspond to the change in Gibbs energy when the solute is transferred from the gas phase to liquid water, which involves a similar change in solute polarization to what occurs during membrane permeation. These nonpolarizable models will have the effect of induced polarization “baked in” through the parameterization, so the PMF barrier will be similar despite the neglect of induced polarization. For example, the permanent dipole moments of molecules described with the nonpolarizable GAFF model are systematically larger than the experimental gas phase values because of a systematic bias in the quantum chemical method used to assign the atomic charges (54; 114). In effect, this method of parameterization mimics the effect of induced polarization, although the physical basis for this is not entirely realistic. Likewise, the accuracy of a polarizable model is only as good as its parameterization.

Polarizable force fields can provide some other advantages for modeling permeation. Because of the larger parameter space and more physically realistic description of the system, they can be parameterized to describe a greater range of properties with greater accuracy. For instance, the SWM4-NDP Drude polarizable water model provides good descriptions of the dielectric constant and diffusion coefficients of water (115). Some popular nonpolarizable water models, like TIP3P, are significantly in error for the calculation of these properties. The water permeation rate calculated by Riahi and Rowley using the Drude model was in quantitative agreement with experiment, suggesting that the solubility-diffusion model combined with a Drude polarizable force field is an effective method for modeling permeation (49). Developing lipid models for use with more accurate nonpolarizable water models, like TIP4P/2005 (116; 117), would address some of these limitations without requiring the use of polarizable models. Future development of both polarizable and nonpolarizable models for membrane permeation would benefit from being validated against the solvation energies and diffusion coefficients of the solutes in water and the liquid alkanes used to parameterize the lipid tails.

There are also several drawbacks associated with the use of polarizable force fields. The number of molecules that have been parameterized for simulations using polarizable force fields still lags behind established nonpolarizable force fields. The computational cost of polarizable models is greater than nonpolarizable force fields due to the need to calculate additional force field terms. Depending on the size and composition of the system, this can range from a factor of 4 for the Drude model to a factor of 20 for the AMOEBA model. Standard dual-Lagrangian implementations of the Drude model are also incompatible with replica exchange methods. As a result, the benefits of using a polarizable model must be weighed against increased statistical error because the configurational sampling is more limited.

Simulations of membrane permeation using polarizable models are now possible for select solutes and lipids. The development of polarizable models for additional solutes and lipids will extend this further in the coming years. These models will make it possible to quantitatively examine the role of polarization in membrane permeation and could allow a greater level of accuracy in cases where induced polarization is significant. The choice between a nonpolarizable or a polarizable model depends on the availability and accuracy of the force fields and whether sufficient sampling can be performed using the polarizable model.

### 5.2. Coarse grain models

Coarse grain models have also been used to simulate membrane permeation. These models reduce the computational expense of the simulation by using a simplified description of intermolecular interactions and grouping atoms into “bead” particles. The MARTINI model for lipids has been notably successful in describing many of the physical properties of bilayers (118; 119). Groups of atoms of a bilayer system are transformed into bead particles. Each bead represents 3–4 non-hydrogen atoms of the lipid or solvent (Fig. 6). The nonbonded interaction potentials of the various types of bead particles are parameterized to reproduce the partitioning of solutes between various solvents (e.g., water–octanol). From this basic design, the surfactant properties of the bilayer arise intrinsically.

**Figure 6:**
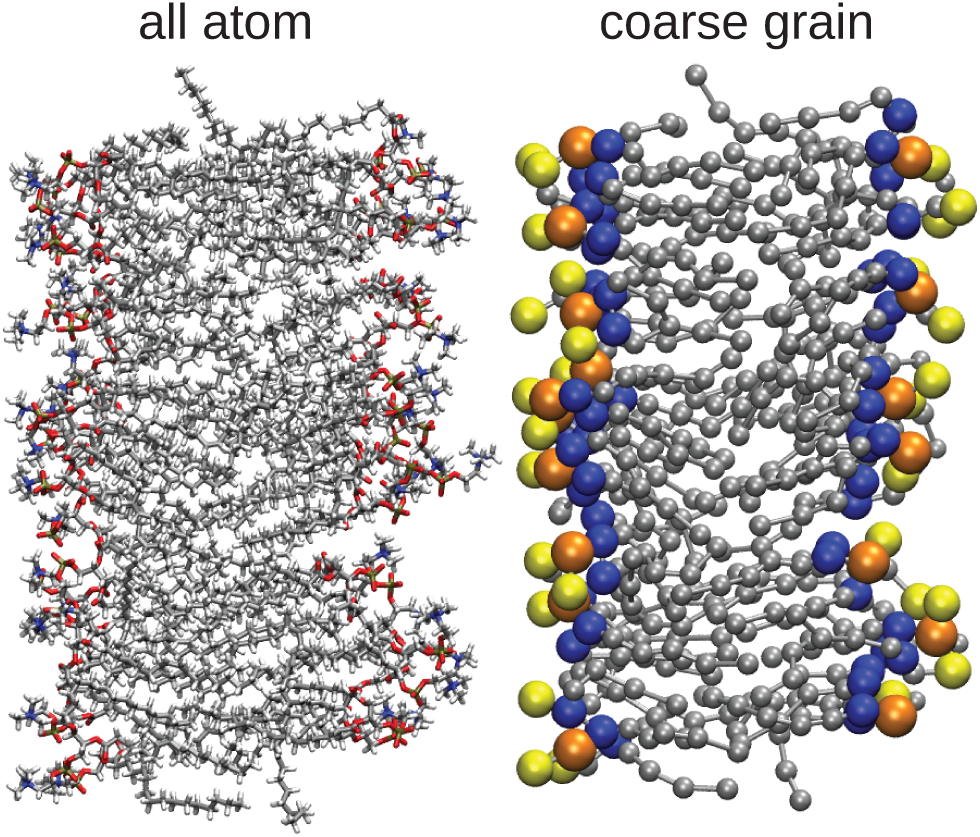
An all-atom POPC model bilayer (left) vs a MARTINI coarse grain model (right). Groups of 3–4 non-hydrogen atoms and the hydrogens bound to them are combined into a single “bead” particle in the coarse grain model.

Through coarse graining, much of the atomic-scale resolution of the system is lost. Despite this, the description of many bilayer physical and dynamic properties relevant to permeation is remarkably good. For example, the viscosity coefficient of MARTINI water is 7 × 10^−4^ Pa·s (at 323 K), in fairly good agreement with the experimental value of 5.5 × 10^−4^ Pa·s (120). The self-diffusion coefficients predicted by this model are more variable; the rate of diffusion of water and alkanes are in good agreement with experiment, but alcohols and lipids diffuse at rates up to 20 times faster than experiment (121). The water permeability of a MARTINI DPPC bilayer was estimated by direct simulation to be 1.5 × 10^−3^ cm/s, which is a reasonable value given the simplicity of the model (118).

Coarse grain models have been used to study many of the same solutes that have been studied using all atom models, including H2O, gases, ions, and small organic molecules. Coarse grain models are particularly suitable for modeling permeation of nano-scale solutes that would be difficult to simulate at an appropriate scale with an atomistic model. These studies have included the permeation and partitioning of fullerenes and nanoparticles. These studies are summarized in Table 1.

**Table 1:**
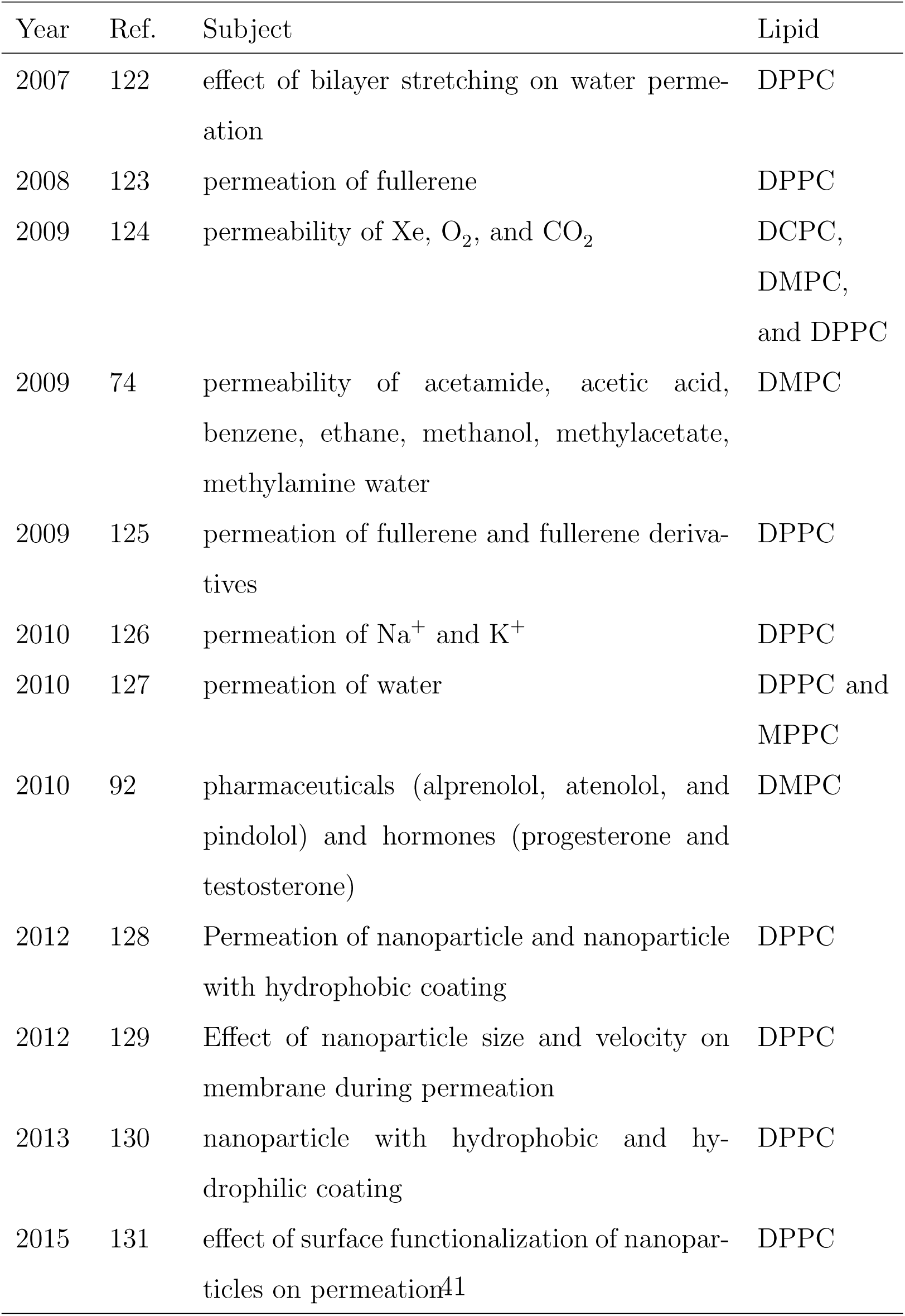
Papers using coarse grain models to simulated membrane permeation.

For molecules that are too small to be coarse grained without losing significant aspects of their chemical properties, multiscale models may be effective. Orsi and Essex developed a model where the permeant was described by an all-atom model but the solution and bilayer were coarse grain (92). This model was generally effective at reproducing the permeability coefficients predicted by all-atom models.

Coarse grain models can be more computationally efficient than all atoms models by multiple orders of magnitude. Consequently, coarse grain simulations of permeation commonly include hundreds of lipids and simulations that are hundreds of nanoseconds long are routine. This is particularly significant when modeling large solutes that require large simulation cells and extensive sampling to calculate accurate *w*(*z*) and *D*(*z*) profiles. This advantage must be weighed against the loss of atomic detail in the model and quality of the coarse grain model used.

## 6. Conclusions

Molecular simulation has contributed significantly to the study of nonfacilitated membrane permeability. These simulations provide atomistic-scale insight into why solutes cross membranes at different rates and why the composition of the membrane affects permeability. The solubility-diffusion model has provided a method for the quantitative calculation of permeability coefficients from the potential of mean force and diffusivity profiles of the solutes on the transmembrane axis. Although accurate calculation of these profiles can be challenging, there are established computational strategies for computing them. More sophisticated molecular mechanical force fields will improve the accuracy of the underlying models. More representative configurational sampling through improvements in computing hardware and more efficient simulation algorithms will reduce statistical error. Coarse grain and polarizable force fields have promise in addressing the sampling issues and accuracy of permeation simulations, respectively. Experimental permeability measurements often rely on complex models and can involve complex bilayer compositions, so direct comparisons to experimental data are not always straightforward. Once these issues are resolved, the quantitative accuracy of the underlying solubility-diffusion model can also be assessed.

There are many interesting subjects in nonfacilitated membrane permeation that remain unexplored. The success of molecular simulation in describing the effect of cholesterol content on permeability suggests that these techniques could be used to understand the effects of other aspects of membrane composition on permeability. Notably, most studies to date have used pure DPPC bilayers as a membrane model and simulations using mixed-lipid bilayers that are more representative of biological cell membranes have not been reported yet. Simulations may also be able to help resolve the controversy over purported exceptions to the Meyer–Overton rule (25; 132; 133; 134; 26).

Cell membranes are remarkable structures that emerge from the amphiphilic properties of the constituent lipids. The selective permeability of lipid bilayers is ultimately a simple phase partitioning effect that arises from complex intermolecular interactions. Appropriately, the solubility-diffusion model is a simple but effective theory for understanding the complex process of permeability. Marrink and Berendsen’s 1994 paper that first demonstrated that permeability coefficients could be calculated quantitatively using molecular dynamics simulations has been cited over 500 times in the last 21 years and is undoubtedly a landmark paper in computational biophysics (43).

## Acknowledgements

We thank NSERC of Canada for funding through the Discovery Grant program (Application 418505-2012). EAW thanks ACEnet and Memorial University for funding. Computational resources were provided by Compute Canada (RAPI: djk-615-ab) through the Calcul Quebec and ACEnet consortia. We thank Dr. Chris Neale and the reviewers for helpful comments.

## Appendix A. Summary of membrane permeability simulations

**Table A.2:**
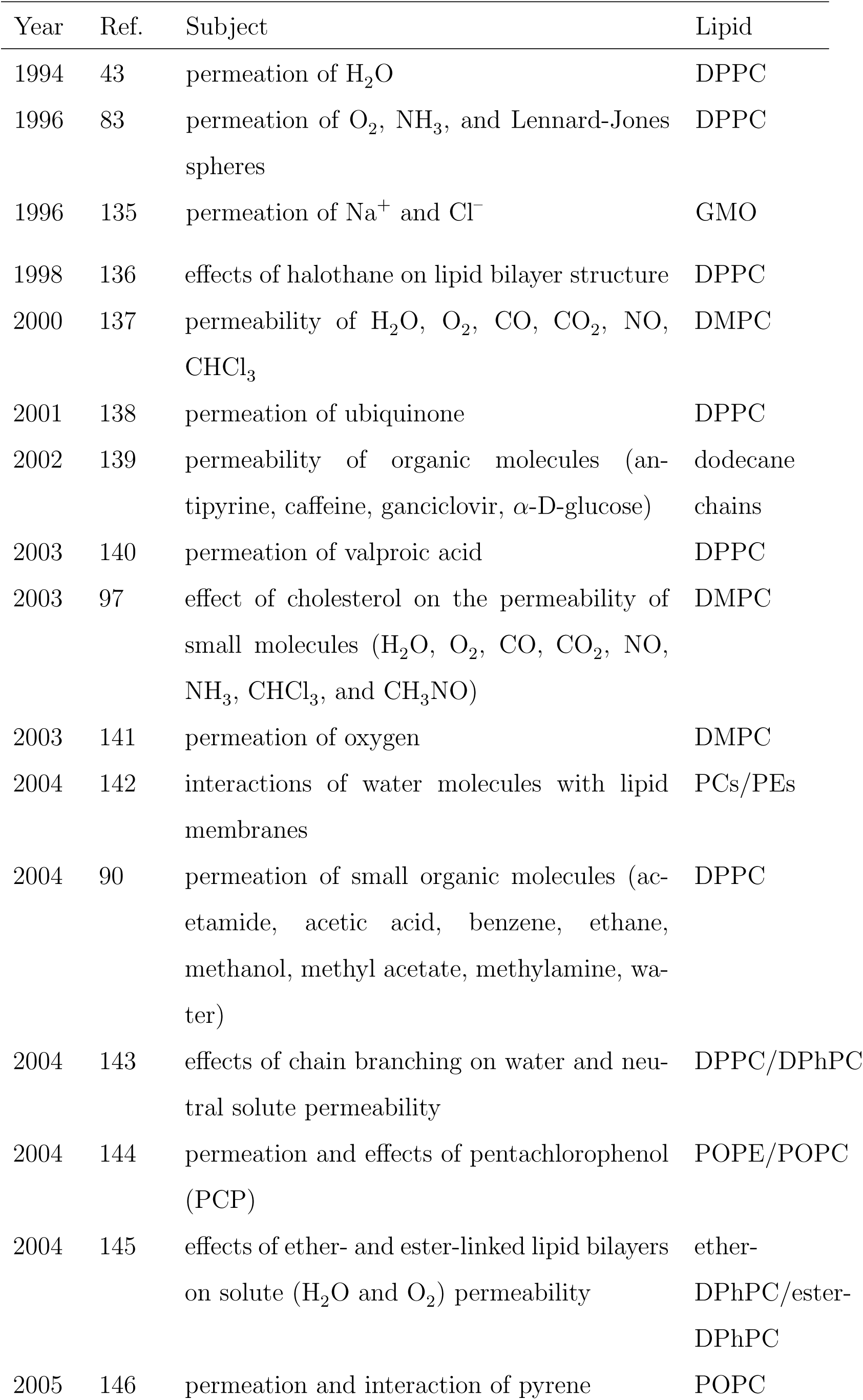
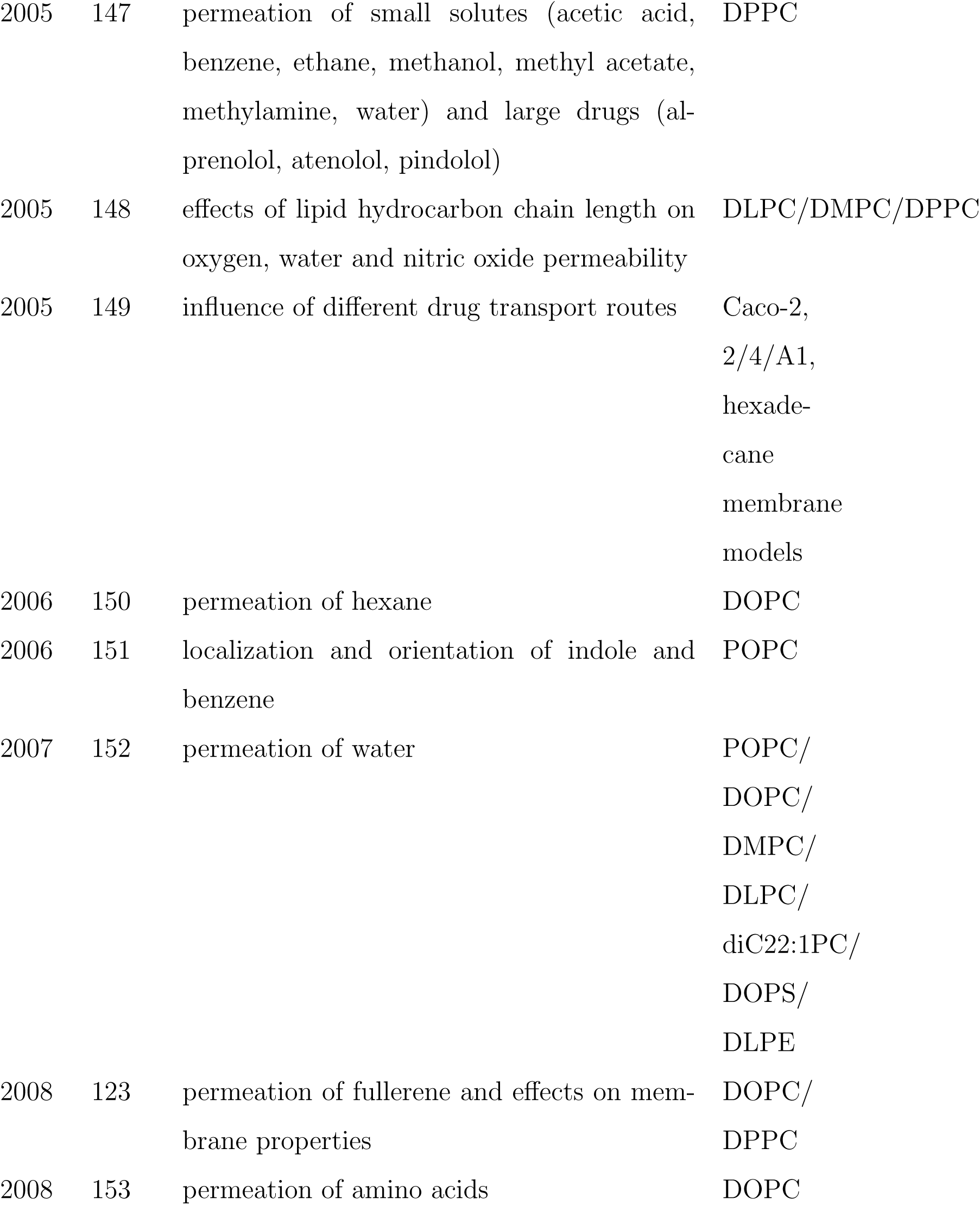
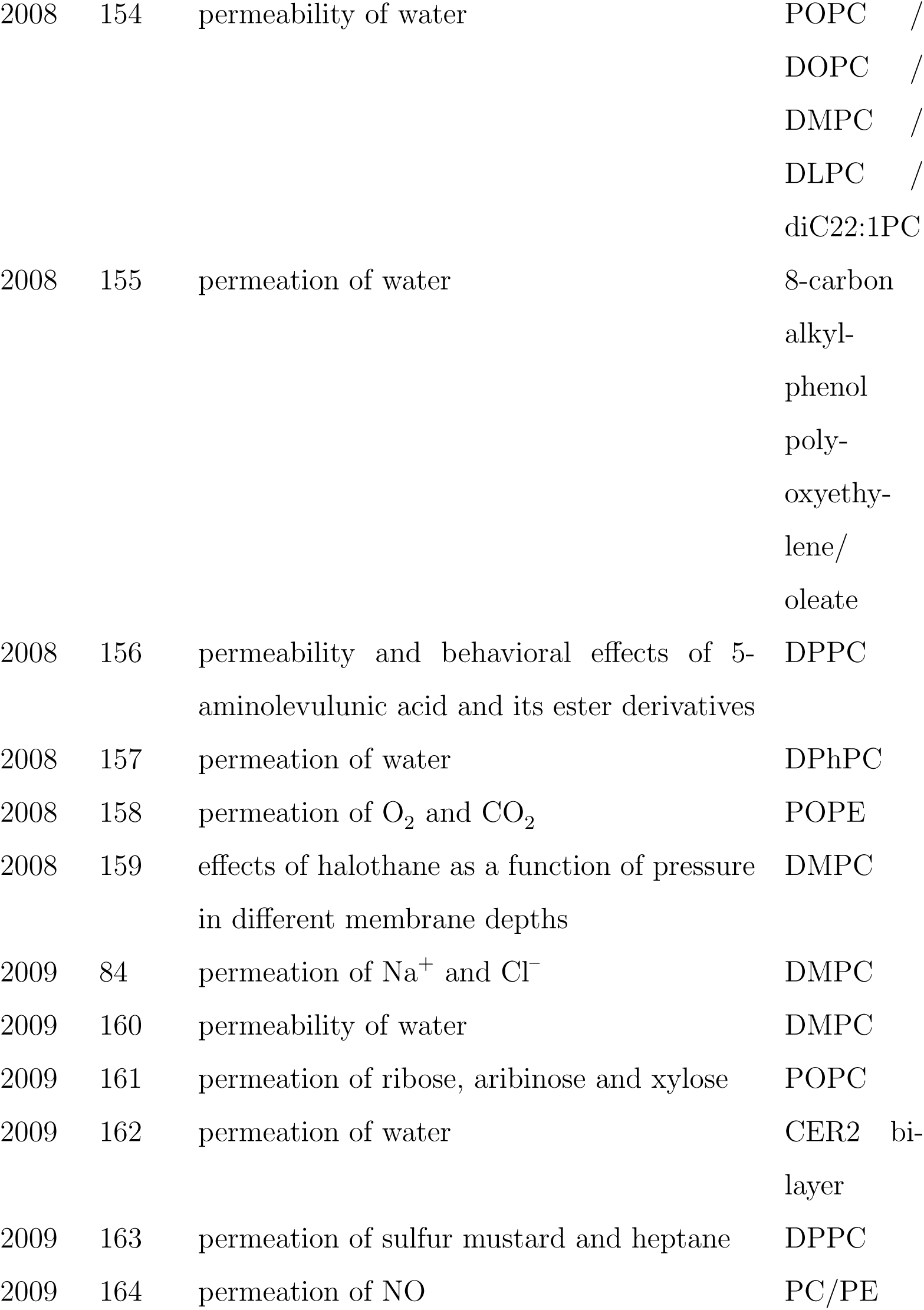
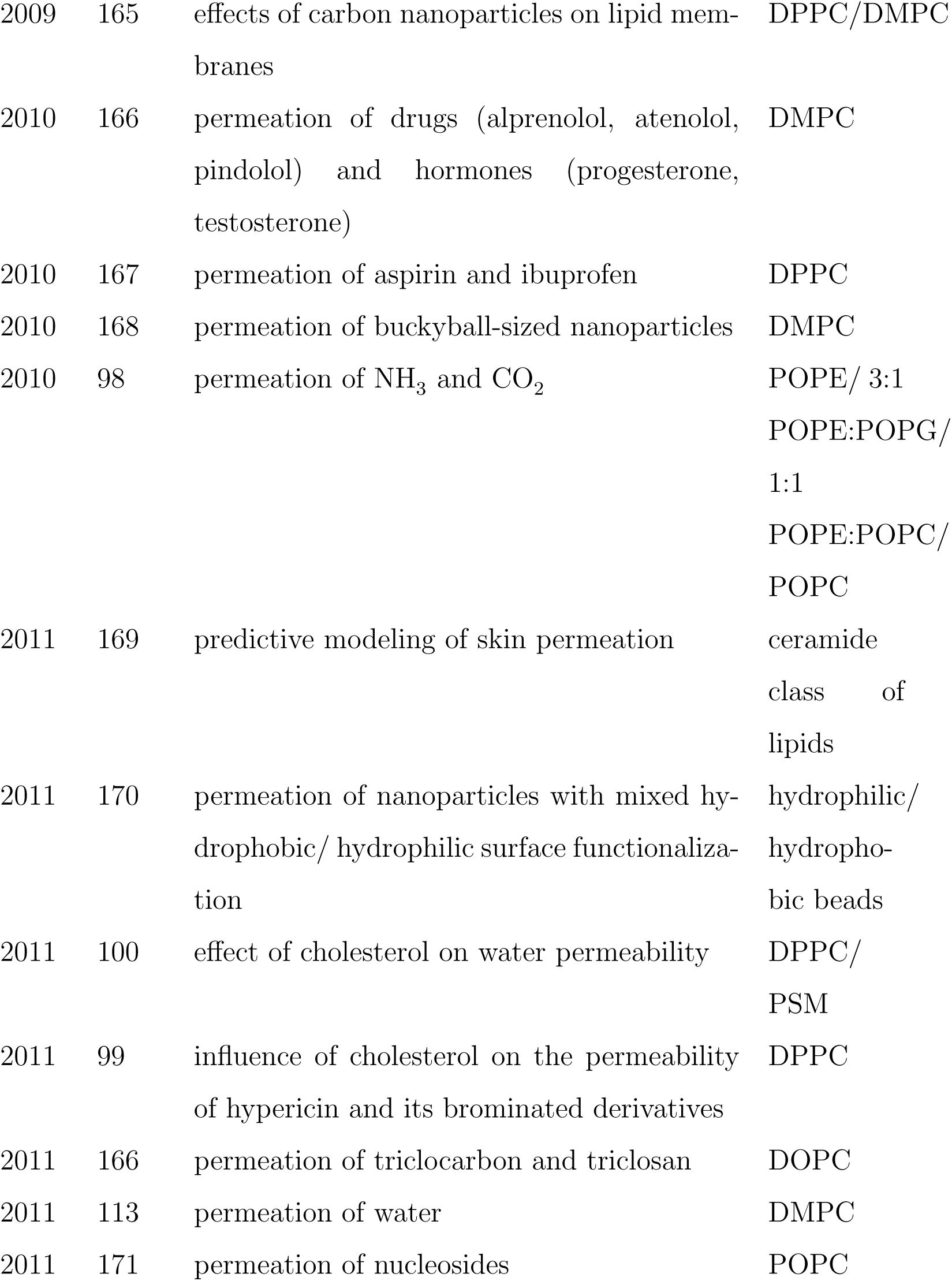
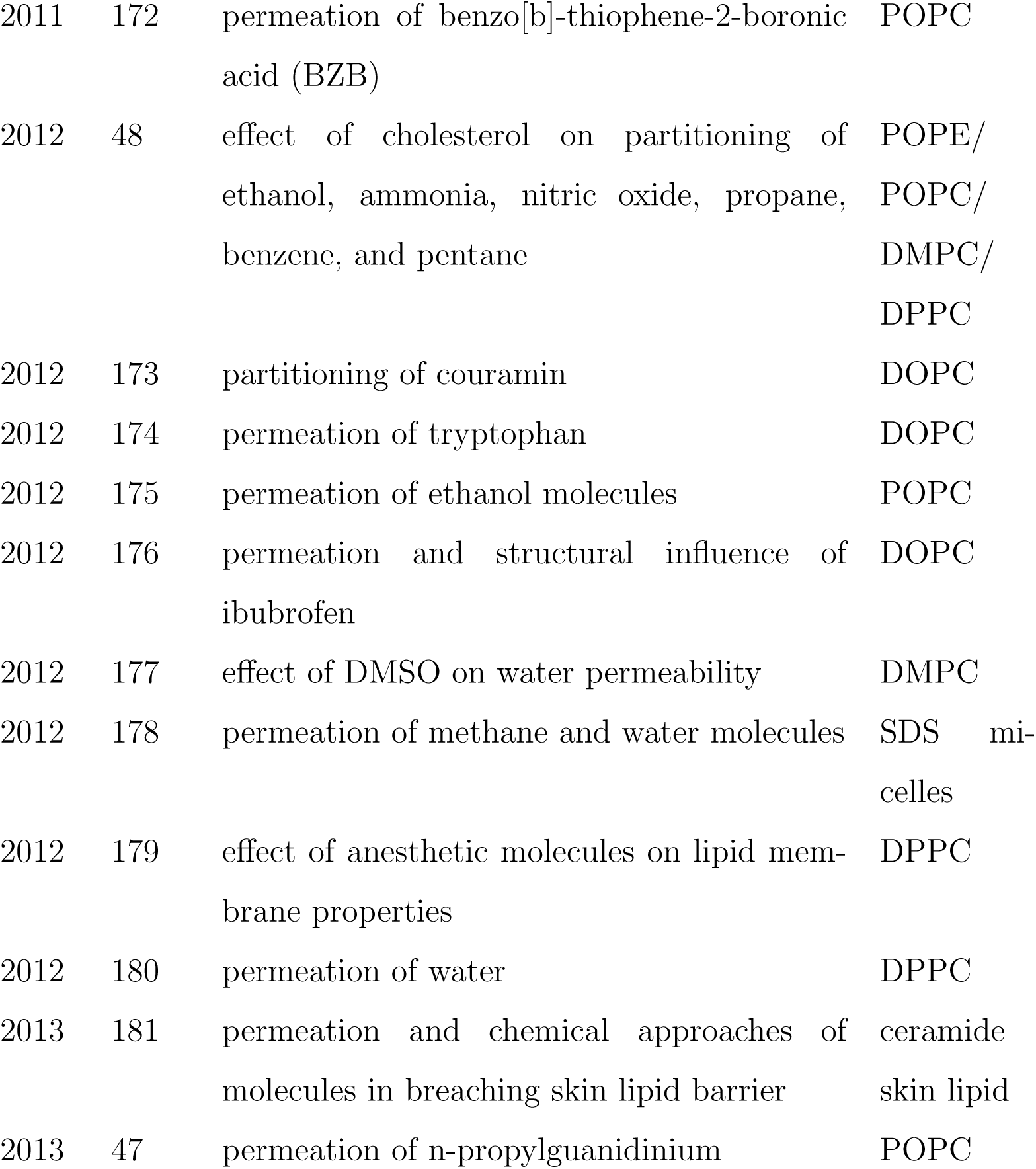
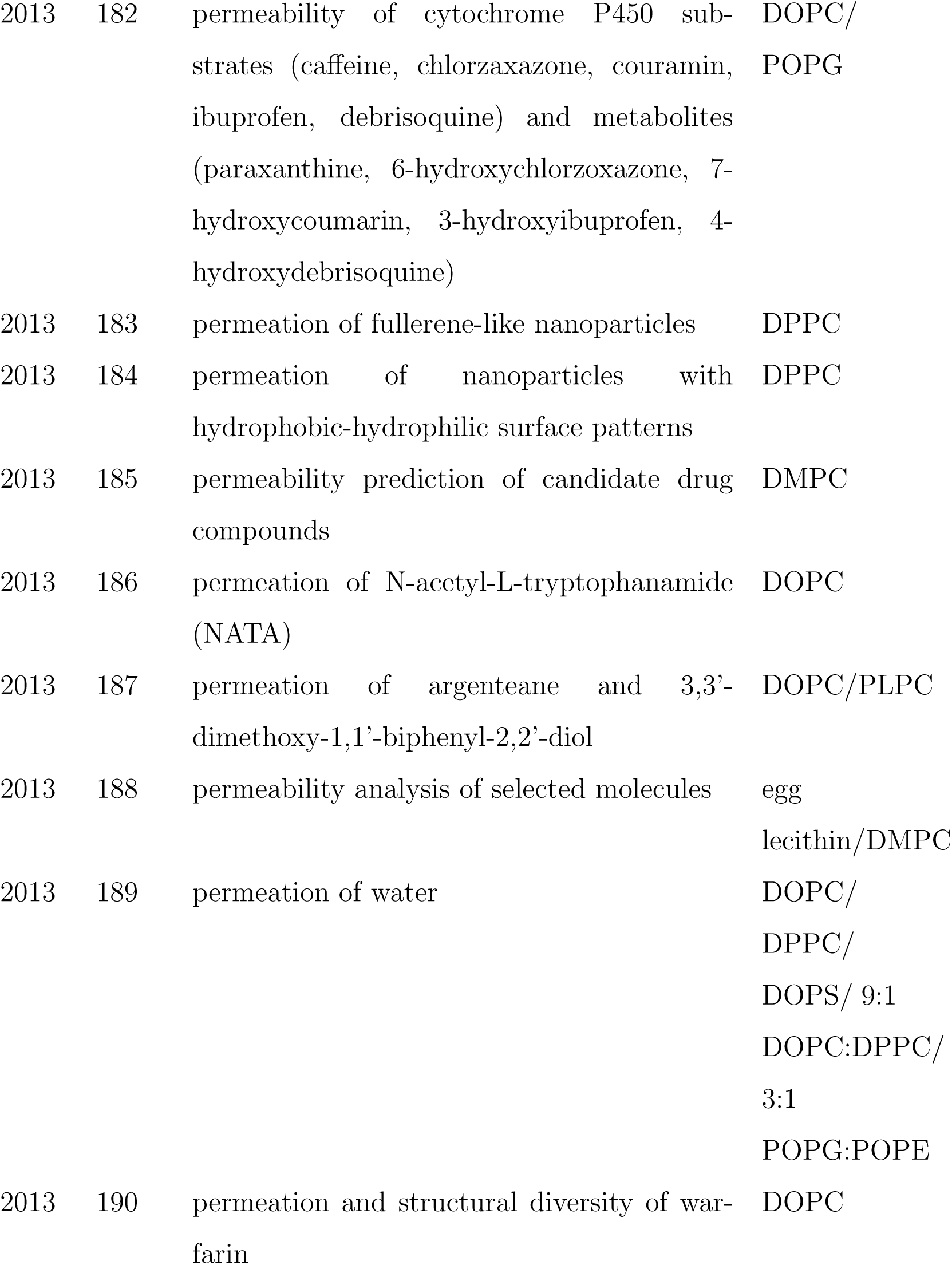
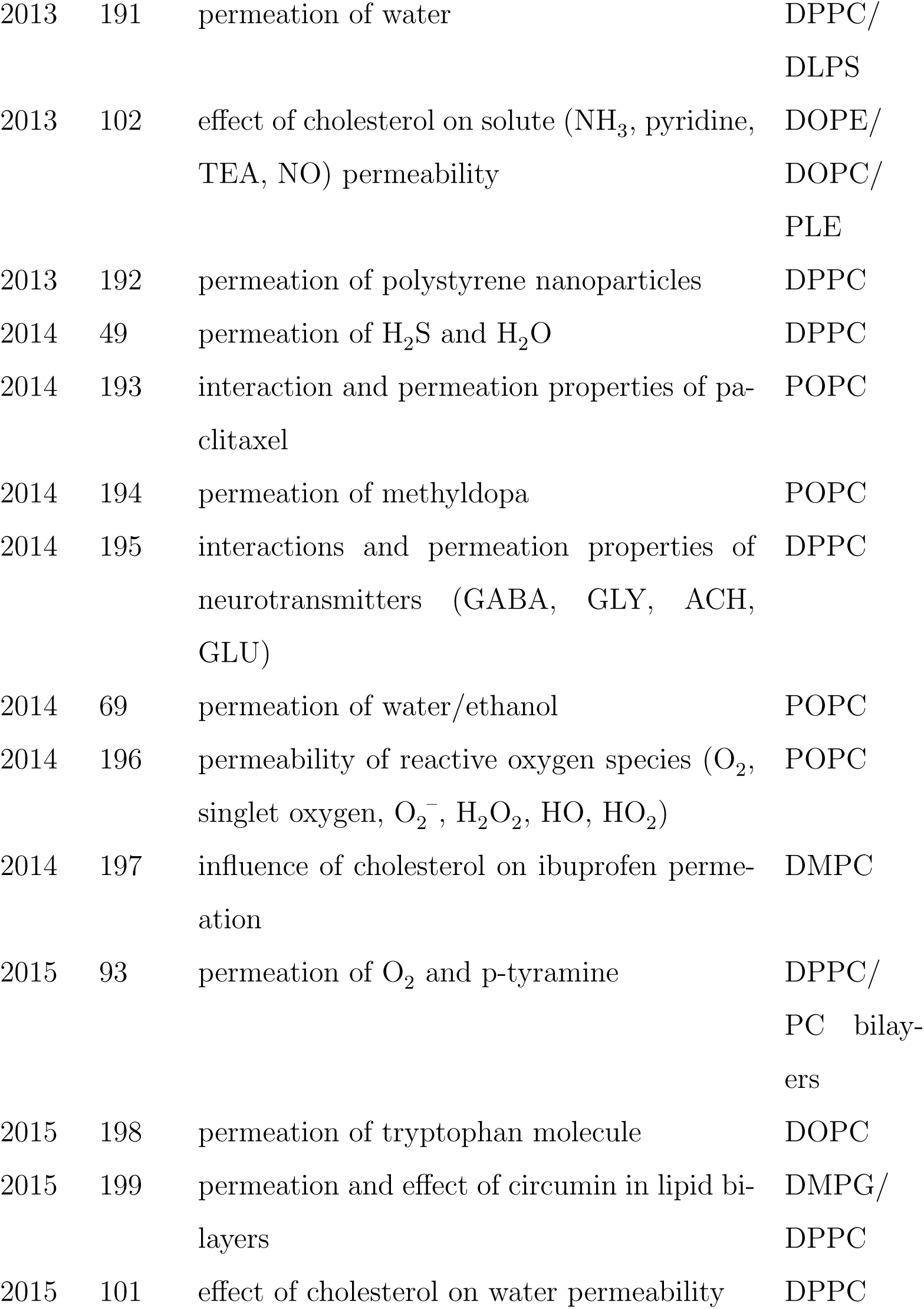
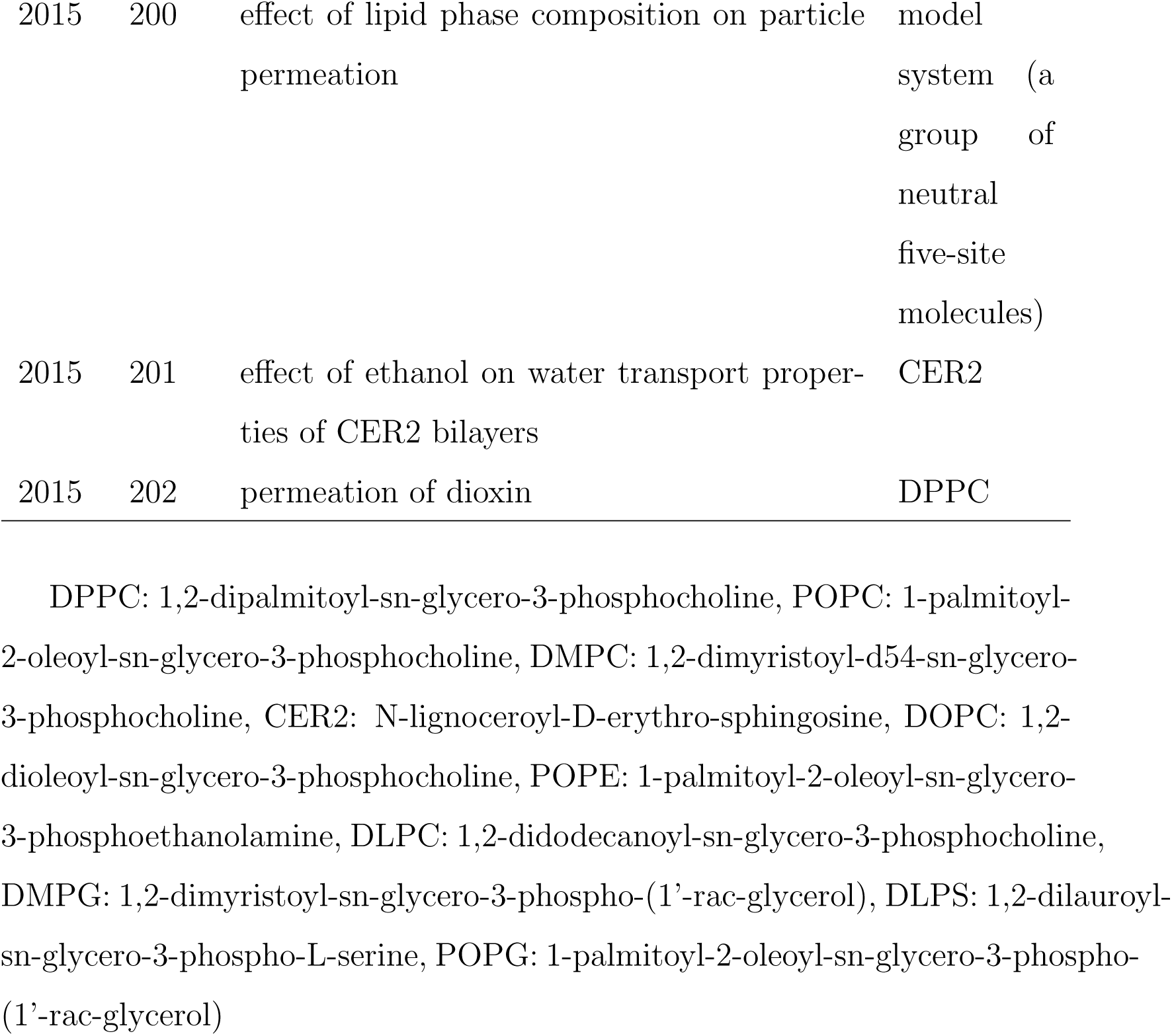
Papers using molecular simulation to model nonfacilitated membrane permeation

## Appendix A. Technical details

The simulations of H_2_O and O_2_ permeation were performed using NAMD 2.10 (203). The CHARMM36 lipid force field was used to represent the DPPC lipids (53). The TIP3P model was used to represent the water molecules (50). The Fischer–Santiago model was used for O_2_. The bilayer model was comprised of 4551 water molecules and 64 DPPC lipids. All bonds containing a hydrogen atom were constrained using the SHAKE algorithm (204). A 2 fs time step was used.

The trajectory of O_2_ permeation was calculated from an equilibriated system where the O_2_ molecule was in solution. An 10 ns NVE simulation was initiated. The 300 ps section where the O_2_ molecule permeates the membrane is presented in Fig. 1.

The PMFs (*w*(*z*)) presented in Fig. 3 were calculated from 20 ns replica exchange umbrella sampling simulation. Each replica corresponded to a NPT simulation of the system where the position of the solute along the *z*-axis was restrained using a harmonic potential (Eqn. 11). The replicas were at 1 Å intervals along the *z* axis over the range *z* = [−40 Å, 40 Å] for a total of 81 replicas. A force constant of *k* = 5 kcal Å^2^ was used for all windows. Exchanges between neighboring replicas were attempted every 1000 time steps. *w*(*z*) was calculated from the probability distributions of these profiles using the weighted histogram analysis method (WHAM) (205; 62). The diffusivity profiles were calculated from the average of six 4 ns MD simulations using Eqn. 17.

